# A minimal model for the role of Rim4 in regulating meiotic exit in budding yeast

**DOI:** 10.1101/2025.08.09.669466

**Authors:** V. Abigail Marquez Davila, Pallavi Gadgil, Anna Zike, Gisela Cairo, Renyu Wang, Sima Setayeshgar, Soni Lacefield

## Abstract

Meiosis ensures formation of haploid gametes through two rounds of chromosome segregation after one round of DNA replication. How this complex cell cycle process is restricted to two and only two divisions is poorly understood. In budding yeast, RNA-binding protein Rim4 binds various mRNAs to prevent their translation. At the onset of meiosis II, phosphorylation and degradation of Rim4, along with the concomitant release of sequestered mRNA, has an important role in ensuring meiotic exit. Building on previous work, we developed a parsimonious mathematical model of meiotic termination that elucidates the role of Rim4-mRNA release and translation of *AMA1* mRNA in the fidelity of meiotic exit. Central to our model is the accumulation of Ama1 protein, a meiosis-specific activator of APC/C. Our mathematical model predicted further outcomes, which we tested experimentally. We found that either slowing Rim4 degradation or disrupting APC/C^Ama1^ activity delayed meiosis II. In some cells, this disruption prevented meiotic exit entirely, leading them to re-enter cell cycle oscillations after meiosis II. These findings demonstrate that the timely activation of this regulatory network is crucial for ensuring irreversible meiotic exit.

## Introduction

The specialized cell division process of meiosis produces haploid gametes from a diploid progenitor cell. This critical reduction in chromosome number is achieved through a single round of DNA replication followed by two distinct rounds of chromosome segregation in which homologous chromosomes segregate in meiosis I and sister chromatids separate in meiosis II. The transition to exit from meiosis is an irreversible step, essential not just for completing cell division, but also for coordinating gamete differentiation (Neiman, 2011; Winter, 2012; MacKenzie and Lacefield, 2020). A dysregulation of meiotic exit has profound consequences, ranging from infertility to the formation of germ cell tumors.

The exit from meiosis requires the halt of the oscillatory activity of cyclin-dependent kinase, Cdk1. The meiotic divisions are controlled by two waves of cyclin-dependent kinase activity, in which Cdk1 is bound and activated by a B-type cyclin (MacKenzie and Lacefield, 2020). During the transition between meiosis I and meiosis II, Cdk1-cyclin B activity declines for anaphase I spindle disassembly but then rises again for metaphase II spindle assembly. The oscillatory activity of Cdk1-cyclin B is controlled through the synthesis and degradation of cyclin B. For both anaphase I and anaphase II onset, Cdk1-cyclin B phosphorylates and activates its inhibitor, the anaphase promoting complex/cyclosome (APC/C), a ubiquitin ligase that when bound to its co-activator Cdc20 ubiquitinates and targets cyclin B for proteasomal degradation. For cells to fully exit meiosis, the cyclin levels and Cdk1-cyclin B activity must remain low.

In budding yeast, after cyclin B levels decline in meiosis II due to APC/C^Cdc20^ activity, a meiosis-specific co-activator of the APC/C, Ama1, binds to APC/C to further ubiquitinate any remaining cyclin B for degradation in anaphase II (Cooper *et al.*, 2000; Cooper and Strich, 2011). APC/C^Ama1^ has other substrates in addition to those of APC/C^Cdc20^, including the middle meiosis transcription factor Ndt80, and other meiotic regulators such as the polo kinase Cdc5 (Cooper *et al.*, 2000; Oelschlaegel *et al.*, 2005; Penkner *et al.*, 2005; Diamond *et al.*, 2009; Tan *et al.*, 2011; Okaz *et al.*, 2012; Argüello-Miranda *et al.*, 2017; Rimal *et al.*, 2020). Ndt80 induces transcription of many genes needed for the meiotic divisions including those that encode the B-type cyclins important for meiosis: *CLB1*, *CLB3*, and *CLB4* (Chu and Herskowitz, 1998; Chu *et al.*, 1998; Hepworth *et al.*, 1998). Therefore, the subsequent proteasomal degradation of cyclins and the transcription factor that induces cyclin gene expression will prevent the accumulation of new cyclins, thereby fully halting the activity of Cdk1, leading to an irreversible meiotic exit.

Several processes temporally regulate the activity of APC/C^Ama1^ during meiosis. First, although the transcription of *AMA1* is upregulated by Ndt80, translation of the mRNA primarily occurs in meiosis II (Chu and Herskowitz, 1998; Brar *et al.*, 2012; Berchowitz *et al.*, 2013; Cheng *et al.*, 2018). *AMA1* mRNA is thought to be sequestered by the translational repressor Rim4, which releases the mRNAs for translation and is subsequently degraded (Berchowitz *et al.*, 2013; Jin *et al.*, 2015; Carpenter *et al.*, 2018; Rojas *et al.*, 2023). The release of mRNA and subsequent degradation of Rim4 are mediated by its phosphorylation state (Carpenter *et al.*, 2018). Second, during meiosis I, APC/C^Ama1^ is inhibited by Polo kinase, Cdc5, specifically when bound to Spo13 (Rojas *et al.*, 2023). Spo13 binds Cdc5 in meiosis I but is then targeted for degradation by APC/C^Cdc20^ at the end of anaphase I. Finally, APC/C^Ama1^ is inhibited by Cdk1-Clb1 (Okaz *et al.*, 2012). Therefore, once *AMA1* mRNAs are released by Rim4 in metaphase II and translated, APC/C^Ama1^ only becomes active after APC/C^Cdc20^ targets Clb1 for degradation. Whether the other B-type cyclins take over this function when Clb1 is absent is currently not known.

Once cells initiate anaphase II, the activities of APC/C^Cdc20^, APC/C^Ama1^, and the meiosis-specific Sps1 kinase are important for anaphase II spindle breakdown and meiotic cytokinesis (Argüello-Miranda *et al.*, 2017; Paulissen *et al.*, 2020; Seitz *et al.*, 2023). The casein kinase Hrr25 promotes meiosis II spindle disassembly by activating the degradation of Clb1 through APC/C^Cdc20^ and Cdc5 through APC/C^Ama1^ (Argüello-Miranda *et al.*, 2017). Sps1 functions downstream of the Hippo-like kinase Cdc15 for the release of the Cdc14 phosphatase from the nucleolus into the cytoplasm (Paulissen *et al.*, 2020). Once released, Cdc14 dephosphorylates substrates of Cdk1-cyclin B (Visintin *et al.*, 1998; Manzano-López and Monje-Casas, 2020). Cytokinesis in budding yeast meiosis is independent of the contractile actino-myosin ring but instead occurs with the closure of the prospore membrane (Taxis *et al.*, 2006; Neiman, 2024). Both APC/C^Ama1^ and Sps1 activity are needed for prospore membrane closure (Diamond *et al.*, 2009; Paulissen *et al.*, 2016).

We sought to further understand the unique meiotic exit regulatory network in budding yeast, focusing on the steps in meiosis II that lead to increased APC/C^Ama1^ activity and the termination of the oscillations of Cdk1-cyclin B. Our previous work showed that the Rim4 translational repressor is degraded by autophagy (Wang *et al.*, 2020). With autophagy inhibition, Rim4 persists, and cells do not exit meiosis. Instead, the autophagy-inhibited cells undergo aberrant rounds of spindle formation, spindle elongation, chromosome segregation, and Cdc14 nucleolar release and return after meiosis II. The mRNAs held by Rim4 are not translated and the substrates of APC/C^Ama1^ are not degraded. Here, our goal was to understand the network that ensures termination of cell cycle oscillations after meiosis II. Using experiments and mathematical modeling, we further define the components of a minimal meiotic exit network. Our model demonstrates how the release of the translational repression of Rim4 mRNA targets, including *AMA1*, allows the accumulation and subsequent activation of APC/C^Ama1^, which is crucial for irreversible meiotic exit.

## Results and Discussion

### Delayed Rim4 clearance results in extra rounds of spindle pole body accumulation and spindle assembly after meiosis II

Rim4 is a key substrate of autophagy, whose degradation has been shown to be important for meiotic exit after meiosis II. Previous work showed that multisite phosphorylation of the Rim4 C-terminus is needed for normal timing of Rim4 clearance (Carpenter *et al.*, 2018). A mutant version of Rim4 with 47 serine and threonine residues in the C-terminus, *rim4-47A*, resulted in delayed degradation of Rim4 and delayed translation of the Rim4 target *CLB3*. We hypothesized that like autophagy inhibition, delayed dissociation of Rim4 from mRNA could also cause a failed meiotic exit, leading to additional rounds of spindle pole body (SPB) and spindle accumulation after meiosis II. In support of this hypothesis, extra SPB foci were reported in the original description of *rim4-47A* (Carpenter *et al.*, 2018), but whether this was due to extra rounds of spindle assembly after meiosis II was not clear. To further test our hypothesis, we performed time-lapse microscopy on wildtype and *rim4-47A* cells. To determine if cells failed to exit meiosis properly, we monitored wildtype and *rim4-47A* cells that expressed Spc42-GFP to monitor spindle pole body (SPB) accumulation and mRuby2-Tub1 to monitor the spindle. After anaphase II and spindle breakdown, wildtype cells exited meiosis normally (Figure 1A). In contrast, 33% of *rim4-47A* cells failed to exit meiosis. Instead, they underwent meiosis II spindle breakdown and then initiated additional rounds of SPB accumulation, spindle assembly, and spindle disassembly (Figure 1B-C). Cells that underwent these additional cell cycle events accumulated between 5-7 SPBs. The variability in the phenotype is likely due to the stochasticity in the network dynamics at the single cell and population levels, as further discussed below (see also Supplementary Information). These results suggest that delayed clearance of Rim4 can lead to a failure in meiotic exit.

**Figure 1.**
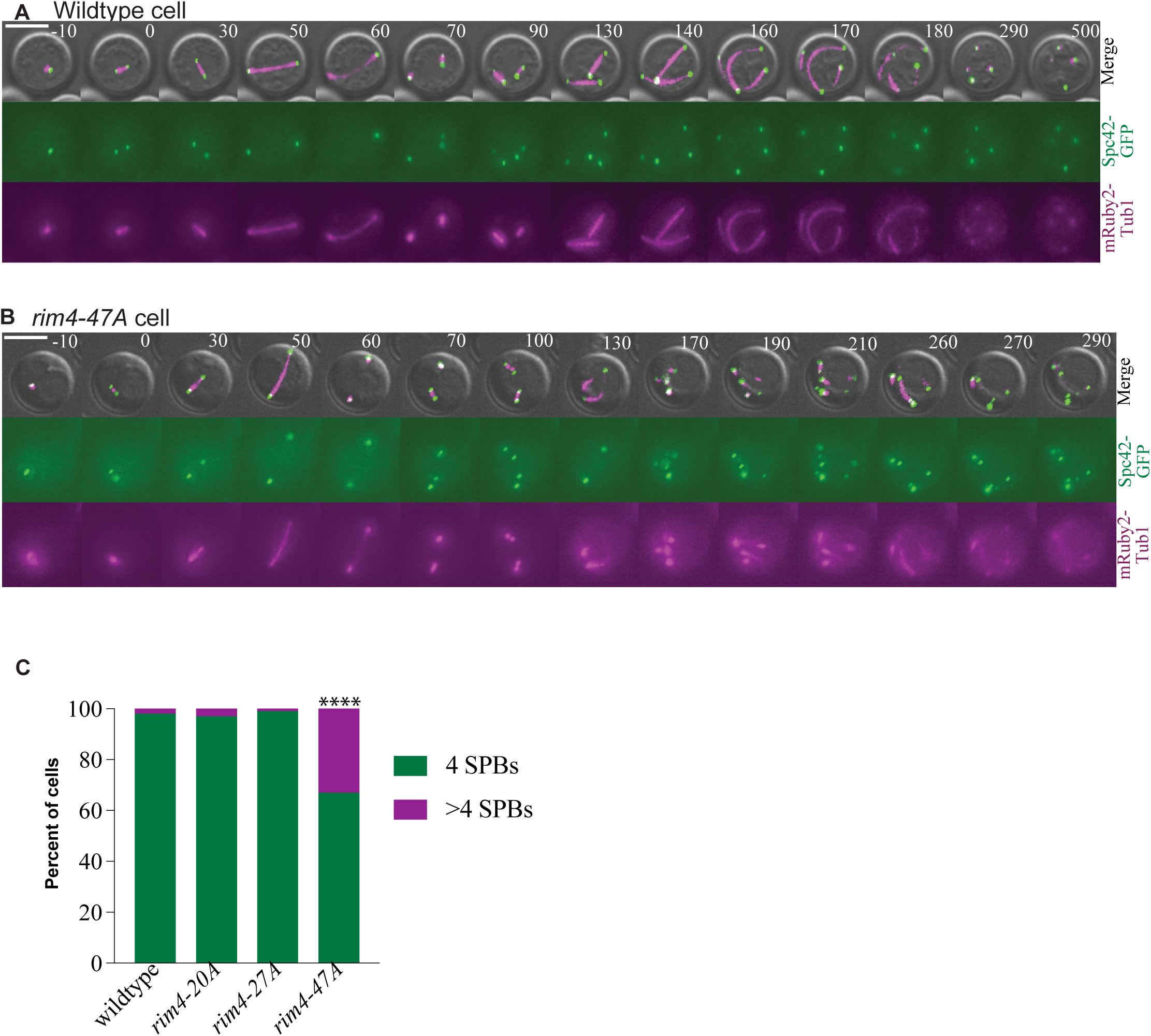
The *rim4-47A* cells undergo extra rounds of SPB accumulation and spindle formation after meiosis II. (A-B) Time-lapse images of a wildtype (A) and a *rim4-47A* (B) cell undergoing meiosis. Cells express *SPC42-GFP* and *mRuby2-TUB1* to monitor SPBs and spindles, respectively. **(C)** Graph showing the percent of cells with 4 or >4 SPBs for each genotype. At least 75 cells counted per genotype. Statistical significance with Fisher’s exact test p <0.0001.

We next tested two other *rim4* phosphorylation mutants, *rim4-27A* and *rim4-20A*. The *rim4-27A* has 27 serine and threonine residues mutated to alanine at the C-terminus from residues 552-718 (Carpenter *et al.*, 2018). The *rim4-20A* has 20 serine and threonine residues mutated to alanine from residues 450-552. Together, the two regions with mutations within *rim4-27A* and *rim4-20A* span the total region of the mutated residues in *rim4-47A* (residues 450-718). Furthermore, *rim4-27A* and *rim4-20A* have delayed Clb3 protein accumulation, but not as delayed as the *rim4-47A* mutants, which suggested that a high level of phosphorylation was needed for Rim4-mRNA release (Carpenter *et al.*, 2018). With time-lapse imaging, we found that almost all *rim4-27A* and *rim4-20A* mutants exited meiosis after meiosis II, similar to wildtype cells (Figure 1C). We conclude that a severe delay in Rim4 degradation, and in translation of Rim4 targets, prevents cells from exiting meiosis after meiosis II.

### The M phase cyclin Clb1, but not Clb3, is important for the normal timing of meiosis II

The rapid degradation of Rim4 in metaphase II suggests a model for a switch-like transition into anaphase II: phosphorylation-mediated dissociation of mRNA-Rim4 aggregates results in both release of the transcripts to activate meiotic exit pathways, as well as targeting of Rim4 for degradation. Of the known Rim4 target transcripts, two have known or potential roles in the regulation of the transition into meiosis II exit: *CLB3* and *AMA1*(Berchowitz *et al.*, 2013; Carpenter *et al.*, 2018). Although Ama1 has an established role in coupling meiotic exit and spore formation (McDonald *et al.*, 2005; Diamond *et al.*, 2009; Argüello-Miranda *et al.*, 2017; Rimal *et al.*, 2020), the role of Clb3 in meiosis II and meiotic exit is unclear. Previous findings demonstrate that Clb3 binds and activates Cdk1 in meiosis II (Carlile and Amon, 2008), suggesting a model that a sudden burst in Clb3 production and Cdk1-Clb3 activity could drive the cells into anaphase II through the phosphorylation and activation of APC/C^Cdc20^. However, other reports have shown that the *CLB3* deletion does not exhibit a defect in sporulation or tetrad formation (Grandin and Reed, 1993; Dahmann and Futcher, 1995). Therefore, we wanted to further investigate whether Clb3 had a more subtle role in meiosis II, such as in maintaining the appropriate timing of meiosis II.

To this end, we performed time-lapse imaging to measure the duration of the meiotic stages and of meiotic exit in wildtype and *clb3Δ* cells. To detect the stages of meiosis, we monitored cells expressing mRuby2-Tub1, which marks the spindle (Figure 2A)(Markus *et al.*, 2015). Metaphase II is measured by the duration from the formation of a bipolar spindle to the time of spindle elongation, which initiates anaphase II (Cairo *et al.*, 2022). Although spindle disassembly can be used as a marker of anaphase II duration, mutants that affect meiotic exit disassemble their spindles by breaking down into fragments. Therefore, scoring the timepoint of full meiosis II spindle disassembly is challenging (Argüello-Miranda *et al.*, 2017; Seitz *et al.*, 2023). Instead, we tagged Cdc14 with GFP and scored the time of Cdc14 release and return to the nucleolus in anaphase II (Figure 2A). When released from the nucleolus, the Cdc14 phosphatase dephosphorylates Cdk1 targets (Visintin *et al.*, 1998). The inactivation of polo kinase, Cdc5, allows the return of Cdc14 to the nucleolus (Visintin *et al.*, 2008). Because Cdc5 is a substrate of APC/C^Ama1^, which becomes active at the end of meiosis II and targets Cdc5 for proteasomal degradation, the return of Cdc14 into the nucleolus can also serve as a proxy to identify the timing of APC/C^Ama1^ activity and the initiation of meiotic exit (Argüello-Miranda *et al.*, 2017).

**Figure 2.**
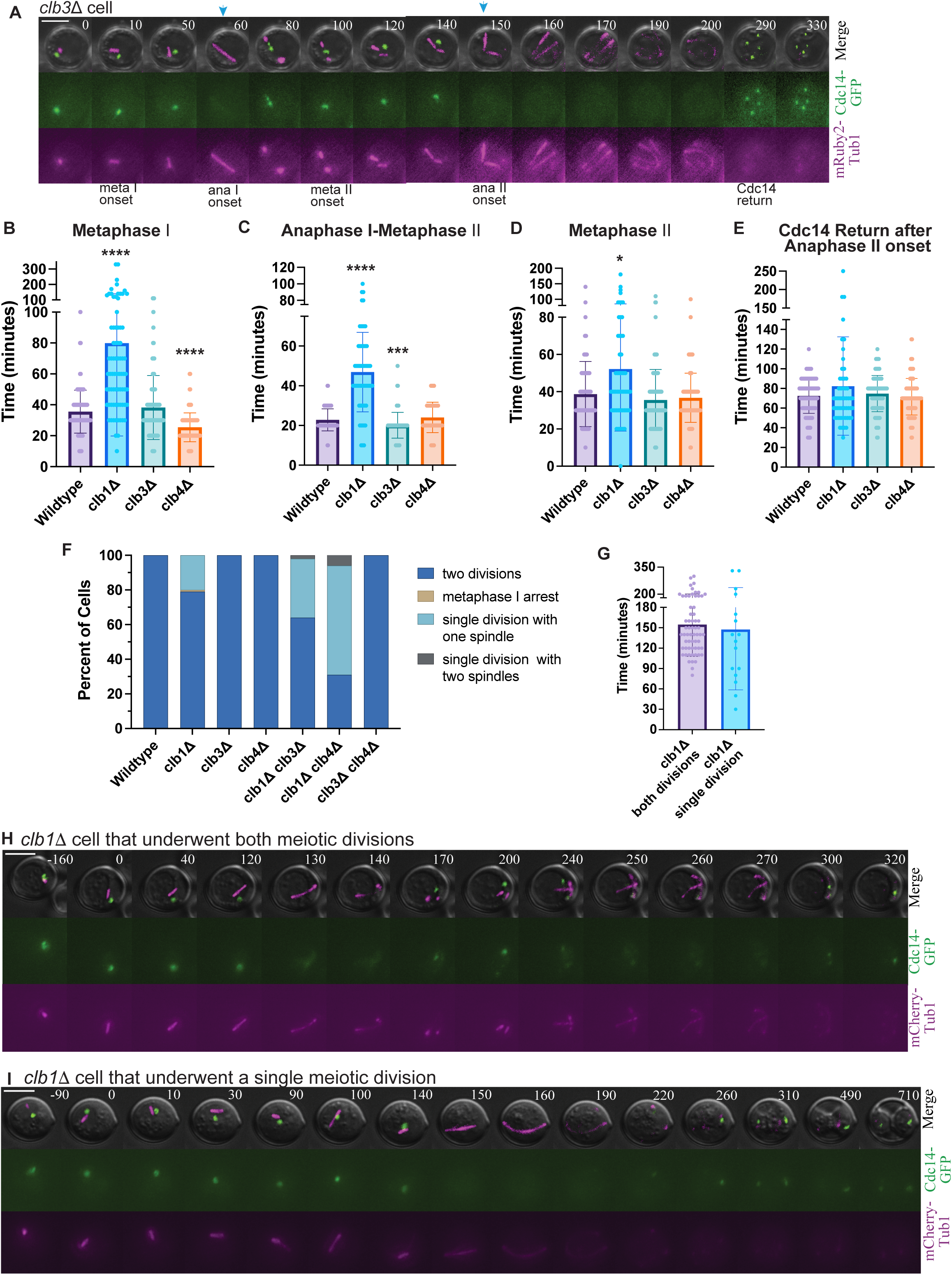
Clb1 is more important than Clb3 and Clb4 in maintaining the normal duration of meiosis II and in ensuring two meiotic divisions. **(A)** Time-lapse images of a *clb3Δ* cell undergoing meiosis. Cells express Cdc14-GFP and *mRuby2-TUB1*. Blue arrows show timepoints of Cdc14 nucleolar release in meiosis I and meiosis II. **(B)** Graph of the mean time from metaphase I spindle formation to spindle elongation at anaphase I. **(C)** Graph of mean time from anaphase I spindle elongation to metaphase II spindle formation. **(D)** Graph of mean time from metaphase II spindle formation to anaphase II spindle elongation. **(E)** Graph of mean time from Cdc14 nucleolar release to Cdc14 return to the nucleolus during anaphase II. **(B-E)** *Indicates statistically significant difference from wildtype (****p < 0.0001, *** p < 0.005, *p<0.05, Mann Whitney test, nΔ 75 cells per genotype). **(F)** Graph comparing the percent of cells that undergo both meiosis I and meiosis II in the single and double *clb* mutant strains. **(G)** Graph comparing the time of both divisions in *clb1Δ* cells that undergo two divisions versus the time of the single division in *clb1Δ* cells that undergo one division. **(H)** Time-lapse images of a *clb1Δ* cell that undergoes two divisions. **(I)** Time-lapse images of a *clb1Δ* cell that undergoes only one meiotic division.

The time of metaphase I, anaphase I-to-metaphase II transition, metaphase II, and Cdc14 return to the nucleolus in anaphase II were similar between wildtype and *clb3Δ* cells (Figure 2B-E). Given that *CLB3* mRNA binds to Rim4 to delay translation to meiosis II (Carlile and Amon, 2008; Berchowitz *et al.*, 2013), we were surprised by these results: we had hypothesized that the absence of Clb3 and the concomitant loss of the surge in Cdk1-Clb3 activity would result in a slower meiosis II. However, based on our experimental results, we conclude that the loss of *CLB3* does not affect the timings of meiosis II or meiotic exit (Figure 2D-E).

Given the partial redundancy of cyclins in the cell cycle, we next investigated whether the other b-type cyclins, Clb1 and Clb4, were important for the normal meiosis II timings. We deleted individual cyclins and monitored meiosis. As previously shown in both W303 and SK1 strain backgrounds, the sporulation of *clb4Δ* cells resulted in an ascus with 4 spores, known as a tetrad. In contrast, the sporulation of *clb1Δ* cells resulted a mixture of cells that underwent one division and two divisions, creating dyad and tetrads, respectively (Dahmann and Futcher, 1995; Rojas *et al.*, 2023) (Figure 2F-I). The live imaging showed that in our W303 background, 79% of *clb1Δ* cells underwent two divisions and the remaining 21% underwent only a single meiotic division (Fig. 2F). Intriguingly, the timing of the singular *clb1Δ* division was similar to the duration of both divisions in *clb1Δ* cells that undergo two divisions, suggesting that the cells delayed in metaphase I undergo only one division, similar to the findings in the SK1 strain background (Rojas *et al.*, 2023) (Figure 2G). As shown previously, the *clb1Δ* cells were also delayed in meiosis I (Carlile and Amon, 2008; Rojas *et al.*, 2023) and in the transition between meiosis I and meiosis II, whereas *clb4Δ* cells were not delayed (Figure 2B-C).

Of those *clb1Δ* cells that undergo two divisions, metaphase II was prolonged by approximately 15 minutes compared to wildtype (Figure 2D). In contrast to *clb1Δ* cells, *clb4Δ* cells had similar timings as wildtype and *clb3Δ* cells in metaphase II. The time of anaphase II was not significantly different in *clb1Δ* and *clb4Δ* cells compared to wildtype. The meiosis II delay in *clb1Δ* cells was unexpected because it was previously thought that Cdk1-Clb1 was not active in meiosis II (Carlile and Amon, 2008). Therefore, Clb1 is needed for the normal timing of meiosis II, but the other M phase cyclins, Clb3 and Clb4, likely compensate for the loss of Cdk1-Clb1 activity.

We wanted to further understand the compensatory mechanisms of the cyclins and used time-lapse microscopy to monitor double mutant combinations of *CLB1*, *CLB3*, and *CLB4*. Previous work showed that *clb1Δ clb3Δ* and *clb1Δ clb4Δ* double mutants had an increase in dyad spores, suggesting a misregulation of meiosis (Dahmann and Futcher, 1995). With live imaging, we found that 36% of the *clb1Δ clb3Δ* double mutant cells that entered meiosis underwent a single meiotic division with a single round of Cdc14 nucleolar release (Figure 2F). Of the double mutants tested, *clb1Δ clb4Δ* mutants had the most severe phenotype with 63% of cells undergoing a single division. In contrast, 100% of the *clb3Δ clb4Δ* double mutants, in which Clb1 is the only M-phase cyclin present, underwent both meiotic divisions. Combined, our results suggest that Clb3 and Clb4 contribute to meiotic regulation, but Clb1 alone is sufficient for ensuring that cells undergo both meiotic divisions.

### A minimal mathematical model for the transition from meiosis II to meiotic exit

Integrating these findings with our previous work and that of others, we can define a minimal network for understanding the role of Rim4 in regulating meiosis II and meiotic exit (Oelschlaegel *et al.*, 2005; Cooper and Strich, 2011; Tan *et al.*, 2011; Okaz *et al.*, 2012; Berchowitz *et al.*, 2013; Argüello-Miranda *et al.*, 2017; Carpenter *et al.*, 2018; Rojas *et al.*, 2023) (Figure 3A). The network is initiated in metaphase II when the Ndt80 transcription factor maintains expression of the APC/C co-activator Cdc20, the M phase cyclins, and the cell cycle kinases Ime2 and Cdc5 (Chu and Herskowitz, 1998; Oelschlaegel *et al.*, 2005; Okaz *et al.*, 2012). Ndt80 also continues to induce its own expression through a positive feedback loop (Chu and Herskowitz, 1998; Winter, 2012). As metaphase II progresses, Rim4 phosphorylation by Ime2 results in dissociation of mRNA from Rim4 and the targeting of Rim4 for autophagic degradation (Berchowitz *et al.*, 2013; Carpenter *et al.*, 2018). The released mRNAs, including *CLB3* and *AMA1,* are then translated, leading to a surge in protein expression.

**Figure 3.**
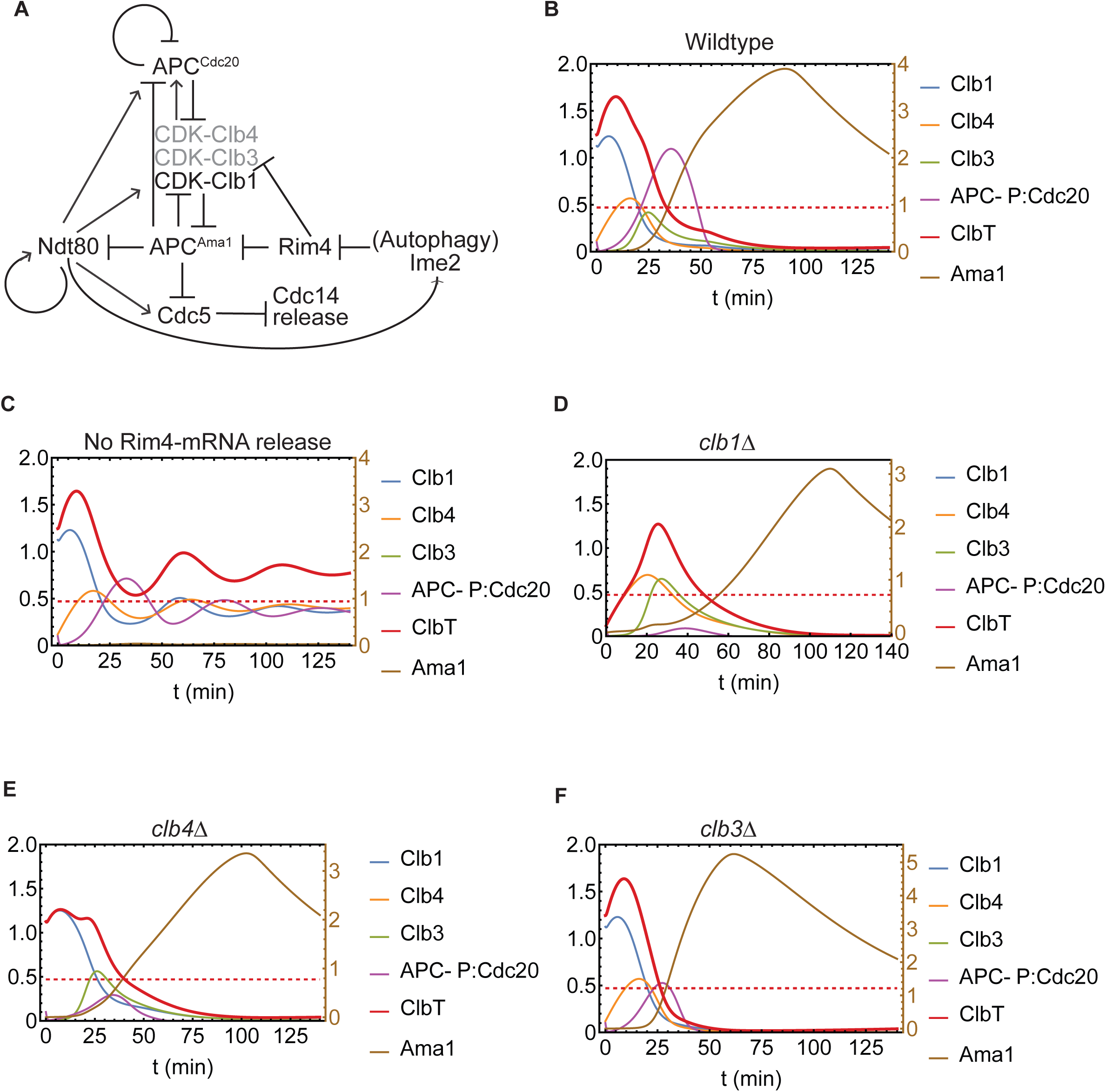
A minimal mathematical model simulates meiotic exit. **(A)** Wiring diagram showing the minimal regulatory network for meiotic exit on which the mathematical model is based. Dynamics for key network elements in **(B)** wild-type cells; **(C)** the absence of mRNA release from Rim4; **(D)** *clb1Δ* mutant; **(E)** *clb4Δ* mutant; **(E)** *clb3Δ* mutant. See Figures S1-S10 for dynamics of all network variables in **(B-E)**, including the variables shown here. **(B-E)** Concentrations are in arbitrary units (a.u.) on the vertical axes. The horizontal axes denote time since the start of meiosis II. The red dashed line indicates a threshold value assumed in the model for total Cdk1-Clb concentration; decline of Cdk1 activity below this threshold is required for proper chromosome segregation and directing the cell toward the sporulation pathway.

The activation of APC/C^Ama1^ requires several regulatory steps that occur as the cell transitions from metaphase II to anaphase II. Previous work has shown that APC/C^Ama1^ activity is inhibited by Cdk1-Clb1, so the presence of Ama1 protein will not immediately result in APC/C^Ama1^ activity (Okaz *et al.*, 2012). We have added Cdk1-Clb4 and Cdk1-Clb3 as minor inhibitors of APC/C^Ama1^ based on our findings that most *clb1Δ* cells can undergo both meiotic divisions, suggesting that Clb1, Clb4, and Clb3 have some compensatory functions (Figure 2C-F).

With increasing Cdk1-Clb activity, APC/C^Cdc20^ is phosphorylated and activated (Rudner and Murray, 2000; Rudner *et al.*, 2000). APC/C^Cdc20^ then targets the cyclins and securin for degradation, resulting in anaphase II onset (Salah and Nasmyth, 2000; MacKenzie and Lacefield, 2020). With decreased Cdk1 activity, APC/C^Ama1^ is activated, leading to meiotic exit by targeting the cyclins, Cdc5, and Ndt80 for degradation (Oelschlaegel *et al.*, 2005; Penkner *et al.*, 2005; Cooper and Strich, 2011; Okaz *et al.*, 2012; Argüello-Miranda *et al.*, 2017). With Ama1 activity, the anaphase II spindle disassembles. The degradation of the Ndt80 transcription factor ultimately results in irreversible meiotic exit because the meiotic regulators are no longer synthesized.

To understand this regulatory network at the systems level and corroborate its predictions with our experiments, we developed a deterministic mathematical model of the wiring diagram, given by a system of nonlinear ordinary differential equations and associated kinetic parameters (See Supplementary Information 1.1-1.3 for model description, equations, limitations, and Tables S1-S10 for governing parameters and initial conditions) (Goldbeter and Koshland, 1981; Goldbeter, 1991; Chen *et al.*, 2000; Tyson and Novak, 2001; Novák and Tyson, 2003, 2008, 2008; Okaz *et al.*, 2012; Qiao *et al.*, 2016; Tyson and Novák, 2022). The model describes the behavior of wildtype cells undergoing meiotic exit during meiosis II (see Supplemental Information; Figure 3B, S1). Given the compensatory role of the B-type cyclins (Clbs), we define the duration of metaphase II as the time for the total cyclin levels to drop below a threshold. A single threshold level, given by approximately one third of the total Cdk1-Clb peak value in wildtype (in arbitrary units), determined by trial and error, can be used to qualitatively explain wildtype and all *clb* mutant phenotypes with respect to meiotic exit (Figure 3B-F, S1-S10, S12). The total cyclin levels initially decline with activation of APC/C^Cdc20^ and then further decline with activation of APC/C^Ama1^. The cyclins do not return after APC/C^Ama1^ activation because the Ndt80 transcription factor is also targeted for degradation.

In our previous experiments, we found that the inhibition of autophagy results in a failure of meiotic exit (Wang *et al.*, 2020). Instead, additional rounds of SPB duplication, spindle elongation, and chromosome segregation ensue. We noted that these oscillations resemble a meiosis II division because chromosome segregation occurred without intervening DNA replication. However, unlike the transition from meiosis I to meiosis II, only a subset of the SPBs re-duplicated. The cells typically increased the number of SPBs one at a time and most cells underwent between 1-3 rounds of SPB accumulation and spindle formation, suggesting that the oscillations decay over time.

In the current model, this behavior is recapitulated as damped oscillations in the total Cdk1-Clb concentrations in the exit network dynamics (Figure 3C, S10). Because the cyclins play redundant roles, we considered the total Cdk1-Clb concentrations in the network dynamics. Previous studies have shown that the negative feedback loop between Cdk1-Clb1 and APC/C^Cdc20^ can lead to oscillations (Milo *et al.*, 2002; Novák and Tyson, 2008; Alon, 2019; Tyson and Novák, 2022). An intermediate step serves to provide a delay between Cdk1-Clb1/4 activation of APC/C^Cdc20^ and APC/C^Cdc20^ inhibition of Cdk1-Clb1/4 (via targeting Clbs for ubiquitination and subsequent degradation). The delay is achieved through Cdk1-Clb1/4 phosphorylation of the APC/C core, which is followed by APC/C activation through binding to Cdc20 (Qiao *et al.*, 2016). Additionally, we allow for APC/C^Cdc20^ self-ubiquitination of Cdc20 for its degradation.

During damped oscillations, after an initial decline in Clbs, the total Clb level - and therefore Cdk1 activity – undergoes cycles of increases and decreases. Consequently, APC/C^Cdc20^ activity also rises and falls. As the total Cdk1-Clb level approaches the threshold in oscillatory dynamics, a new cell cycle can be initiated. However, if the total Cdk1-Clb level remains above or below the threshold, the differences between high and low Cdk1-Clb activity may not be sufficient to allow additional rounds of chromosome segregation.

We next tested our model by analyzing the behavior of deletion of *CLB1*, *CLB3,* and *CLB4* genes individually (Figure 3C-E, S1-S7). We then compared the observed phenotype of these strains with the results of the simulated network dynamics (Figure 2C-E, 3D-F). In the *clb1Δ* cells that undergo both divisions, the model recapitulates the observed delay in meiosis II (Figure 2C-E, 3D, S2). For the *clb4Δ* cells, the normal timing of meiosis II that we observed in the experiments agrees with the computed behavior (Figure 2C-E, 3E, S3). In the model, *clb3Δ* cells undergo meiosis II somewhat faster, but within the range of the experimental parameters (Figure 2C-E, 3F, S4). Therefore, our mathematical model simulates the experimental observations for the cyclin mutants.

In this work, we have adopted a deterministic model and numerical simulation approach to describe the regulatory network in Figure 3A. However, we note that the inherent stochasticity arising from thermal energy of biomolecules gives rise to intrinsic fluctuations in the concentrations of components and rates of reactions constituting biological circuits, thereby constraining the accuracy of biochemical processes (Elowitz *et al.*, 2002; Swain *et al.*, 2002; Ghaemmaghami *et al.*, 2003; Bialek and Setayeshgar, 2005; Raj and Van Oudenaarden, 2008; Brar *et al.*, 2012; Thattai, 2016). A major source of intrinsic noise can be attributed to low copy numbers of mRNA species in gene-protein regulatory networks (Newman *et al.*, 2006). While past theoretical work points to the universality of Poisson statistics for mRNA dynamics, the resulting distribution of fluctuating protein concentrations depends on many factors, including their function and expression level (Newman *et al.*, 2006; Thattai, 2016). Measured coefficient of variation (given by standard deviation/mean) for protein numbers can be significant, varying between 10-40% in yeast (Newman *et al.*, 2006). To address the role of noise in the exit network presented here, a fully or hybrid stochastic implementation is the subject of future work (Gillespie, 1977, 2007; Barik *et al.*, 2010, 2016; Ahmadian *et al.*, 2020). In particular, we expect that the decline, followed by threshold crossing of noisy Cdk1-Clb levels in stochastic simulations will give rise to variability in cell fate outcomes and timing of exit that can be compared with the experimentally observed distributions. For cells exhibiting oscillatory dynamics (Figure 3C, 4B, 5A), we expect that stochasticity in protein numbers as well as in the biochemical regulation of phosphorylation-mediated mRNA-Rim4 dissociation and Rim4 clearance by autophagy will lead to population-level heterogeneity in the additional rounds of divisions (4SPBs or >4SPBs).

**Figure 4.**
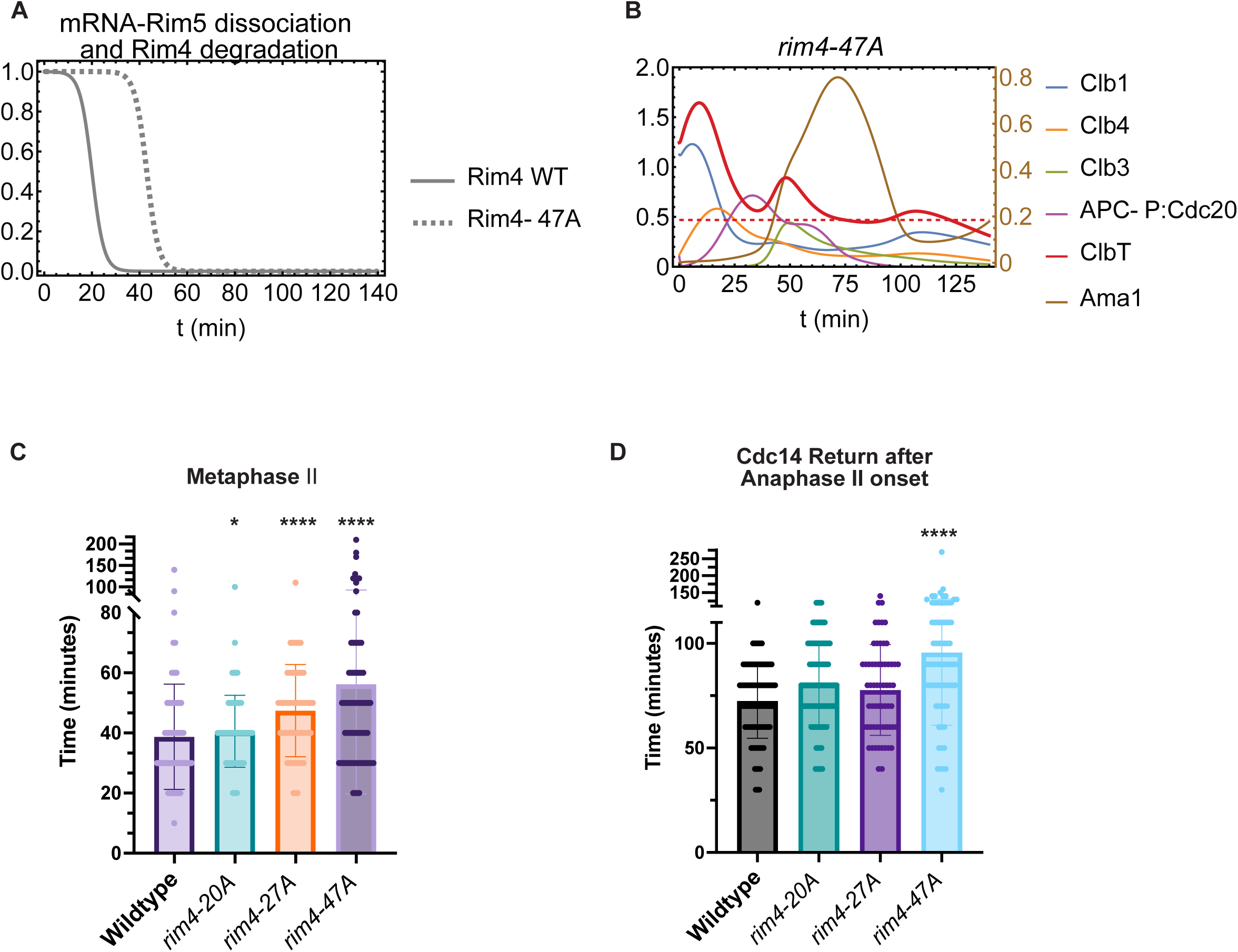
Delaying mRNA-Rim4 dissociation delays meiosis II and results in additional oscillations after anaphase II. **(A)** Dynamics of mRNA-Rim4 complex dissociation in wild-type and Rim4 mutant, demonstrating the model assumptions: (i) multisite phosphorylation of Rim4 results in a sigmoidal, or switch-like, dissociation of Rim4-mRNA complex, and resulting mRNA release for translation, and (ii) mutations in Rim4 phosphorylation sites result in delayed release of mRNA (see also Figure S11). **(B)** Network dynamics when mRNA release from Rim4 is delayed, demonstrating damped oscillations in total Cdk1 activity (see also Figure S12). **(A-B)** Concentrations are in arbitrary units (a.u.) on the vertical axes. The horizontal axes denote time since the start of meiosis II. **(C)** Graph of the mean time from metaphase II spindle formation to anaphase II spindle elongation in different *rim4* mutant strains. **(D)** Graph of mean time from Cdc14 nucleolar release to Cdc14 return to the nucleolus during anaphase II in *rim4* mutant strains. *Indicates statistically significant difference from wildtype (****p < 0.0001, ** p < 0.007, *p<0.05, Mann-Whitney test, nΔ 75 cells per genotype).

**Figure 5.**
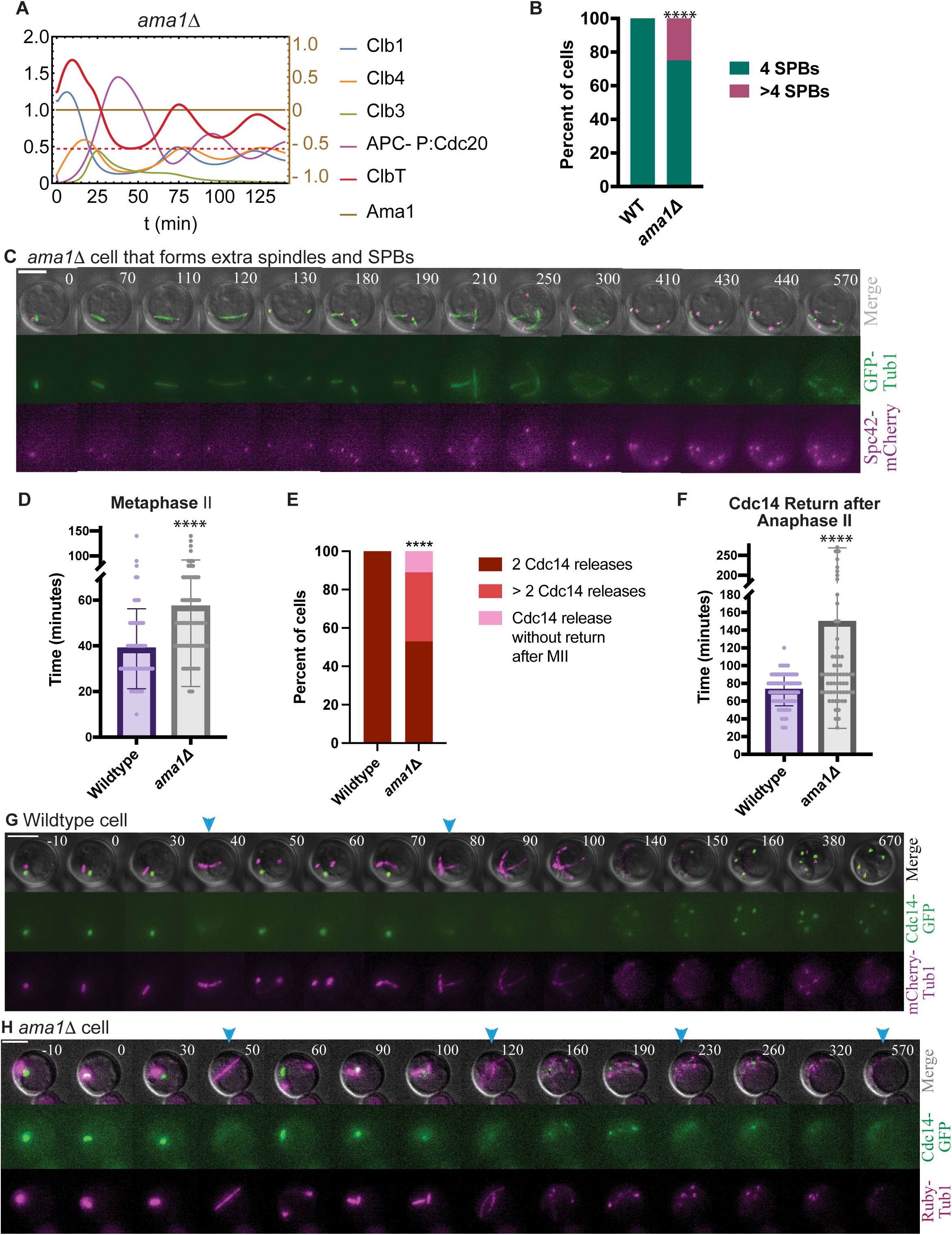
Loss of *AMA1* delays meiosis II and results in additional oscillations after anaphase II. **(A)** Network dynamics for the *ama1Δ* mutant, demonstrating damped oscillations in total Cdk1 activity analogous to mutants in which mRNA release from Rim4 is delayed or suppressed. Concentrations are in arbitrary units (a.u.) on the vertical axes. The horizontal axes denote time since the start of meiosis II. **(B)** Graph showing the percent of cells with 4 or >4 SPBs for each genotype. At least 100 cells counted per genotype. *Indicates statistically significant difference from wildtype (Fisher’s exact test, **** p <0.0001). **(C)** Time-lapse images of an *ama1Δ* cell undergoing meiosis. Cells express *SPC42-mCherry* and *GFP-TUB1* to monitor SPBs and spindles, respectively. **(D)** Graph of the mean time from metaphase II spindle formation to anaphase II spindle elongation. *Indicates statistically significant difference from wildtype (****p < 0.0001, unpaired t test with Welch’s correction, nΔ 75 cells per genotype) **(E)** Graph showing the percent of cells with 2 or >2 Cdc14 releases from the nucleolus. At least 100 cells counted per genotype. *Indicates statistically significant difference from wildtype (Fisher’s exact test, **** p <0.0001). **(F)** Graph of mean time from Cdc14 nucleolar release to Cdc14 return to the nucleolus during anaphase II in *rim4* mutant strains. *Indicates statistically significant difference from wildtype (****p < 0.0001, unpaired t test with Welch’s correction, nΔ 75 cells per genotype). **(G-H)** Time-lapse images of a wildtype cell (G) and an *ama1Δ* cell (H) undergoing meiosis. Cells express Cdc14-GFP and *mRuby2-TUB1*. Blue arrows show timepoints of Cdc14 nucleolar release in meiosis I and meiosis II.

### Testing predictions of the model: delayed mRNA-Rim4 dissociation and Rim4 degradation causes a delayed meiosis II and Cdc14 return

We next used the mathematical model to guide our experimental analysis. Specifically, we investigated the effect of the timing of Rim4-mRNA aggregate dissociation and Rim4 clearance on meiotic exit. Due to absence of experimental details, we do not model the phosphorylation mediated dissociation of aggregates and subsequent Rim4 decay. Rather, we assume a phenomenological time dependence, consistent with regulation via multi-site phosphorylation cascades (see Supplemental Information section 1.1.3). We find that if the onset of sigmoidal decline is strongly delayed with respect to wildtype, as it is in *rim4-47A* cells, the timing of meiosis II is delayed, and the total cyclin levels initially decline, but then reaccumulate for a possible extra cell cycle (Figure 4A-B). These modeling results are consistent with our experimental data describing extra cell cycle oscillations in *rim4-47A* cells (Figure 1B-C).

To further test this prediction experimentally, we analyzed the timing of meiosis II in the strains with mutations in Rim4 phosphorylation sites. Previous work showed that the *rim4-47A* was delayed in meiosis II compared to wildtype cells, but did not show the timing of each stage (Carpenter *et al.*, 2018). With time-lapse imaging, we measured the duration of metaphase II and anaphase II. We found that *rim4-20A* had similar timings as wildtype cells (Figure 4C-D). In contrast, in *rim4-27A* mutants, metaphase II was delayed by an average of 17 minutes, but anaphase II was not delayed. The *rim4-47A* cells were delayed by approximately 20 minutes in both metaphase II and anaphase II. These results, combined with those from Figure 1, support the predictions from the model that delaying Rim4-mRNA release and subsequent translation slows meiotic exit and allows cell cycle oscillations to continue after meiosis II (Figure S12).

We describe the dissociation of mRNA-Rim4 complexes and Rim4 clearance phenomenologically, given by a single, sigmoidal function of time (Figure 4A, Table S11). Cooperative models of multisite phosphorylation kinetics applied to Rim4 amyloid aggregates could plausibly describe the transition from the mRNA-bound (and unphosphorylated) to the mRNA-unbound (and phosphorylated) state (Monod *et al.*, 1965; Koshland *et al.*, 1966; Qian, 2003; Bialek and Setayeshgar, 2008; Salazar and Höfer, 2009; Marzen *et al.*, 2013; Ferrell and Ha, 2014a, 2014b, 2014c). These models display similar sigmoidal functional behavior as the phosphorylation level of Rim4 increases with time, motivating our description. Finally, as also shown in Fig. 4A, we assume that for mutant versions of Rim4 resulting in its reduced phosphorylation, the onset of dissociation of mRNA-Rim4 complex and Rim4 clearance is delayed while the rate of dissociation/clearance (slope of sigmoidal transition) is approximately unchanged. These assumptions are plausibly achieved within mechanistic models of cooperativity.

Finally, we also note that degradation of Rim4 is relevant for the dynamics of the exit network only in so far as ensuring that released mRNA is not re-sequestered. Even if mRNA does not rebind to phosphorylated Rim4, phosphatase activity could result in sufficient dephosphorylation of Rim4 for mRNA rebinding to occur. The present model assumes no rebinding of mRNA (or equivalently, rapid clearance of Rim4). Increasing the delay in the release of mRNA (and clearance of Rim4) leads to later Cdk1-Clb threshold crossing times for Rim4 mutant strains and eventual transition to oscillatory dynamics (Figure S12). Table S11 provides the threshold crossing times for Rim4 and other mutant strains from numerical simulations, in agreement with the experimentally measured mean durations of metaphase II.

### Testing predictions of the model: loss of Ama1 causes additional rounds of SPB accumulation and spindle formation

Finally, we used the mathematical model to predict how the loss of APC/C^Ama1^ activity would affect the termination of oscillations after meiosis II. We eliminated Ama1 from the model to simulate the loss of APC/C^Ama1^ activity and found that our model predicts that oscillations in total cyclin levels occur, leading to oscillations in Cdk1-Clb activity (Figure 5A, S8). Whether cells underwent additional cell cycle oscillations would depend on the total cyclin levels decreasing below and then arising above a threshold. However, if the oscillations in cyclin levels are not proximal to the threshold but remain above or below it, additional cell cycle events would not occur.

We tested these predictions of the model with experiments. By monitoring Spc42-GFP and Tub1-mRuby2, we found that 25% of *ama1Δ* cells increased the number of spindles and SPBs after meiosis II (Figure 5B-C). As previously mentioned, *ama1Δ* cells fragment their spindles during spindle breakdown, making it difficult to assess whether a new shorter spindle assembled (Argüello-Miranda *et al.*, 2017; Seitz *et al.*, 2023). However, we noticed the acquisition of additional SPBs and short spindles that formed between two SPBs, suggesting that we were observing real spindles and not spindle fragments (Figure 5C). On average, metaphase II was delayed by approximately 20 minutes (Figure 5D).

In wildtype cells, after Cdc14 is released from the nucleolus in meiosis II, it returns to the nucleolus approximately 70 minutes later (Figure 5E-G). In contrast, Cdc14 did not return to the nucleolus in 11% of *ama1Δ* cells (Figure 5E). Cdc14 underwent multiple releases in 36% of *ama1Δ* cells (Figure 5E, H). Furthermore, in the cells in which Cdc14 did return in anaphase II, the timing was delayed on average 80 minutes compared to wildtype (Figure 5F). These results suggest that *ama1Δ* cells can undergo additional cell cycle oscillations, as the model predicted.

## CONCLUSIONS

Building on previous findings, we have used mathematical modeling and experimental analysis to define a minimal network for meiosis II exit. Central to our model is the ultimate activation of APC/C^Ama1^ to allow for the degradation of the network components required for cell cycle oscillations. Our model describes a complex pathway involving the release of mRNA from Rim4 aggregates and subsequent degradation of the Rim4 translational repressor and release of the *AMA1* mRNA for translation. With the accumulation of Ama1 protein, APC/C^Ama1^ activity is initially inhibited in meiosis II by Cdk1-Clb1 and Cdk1-Clb4. Once APC/C^Cdc20^ becomes active and targets the cyclins for ubiquitination and subsequent proteasomal degradation, APC/C^Ama1^ is activated and then further targets the cyclins, Ndt80, and Cdc5 for proteasomal degradation. Finally, our mathematical model predicted further outcomes that we then tested experimentally to find that with (i) delayed or suppressed release of mRNA from Rim4 and Rim4 clearance or (ii) loss of APC/C^Ama1^ activity, meiosis II is delayed, and some cells have a failure in meiotic exit, such that cell cycle oscillations occur after meiosis II. Our results demonstrate that the activation of this minimal regulatory network is critical for the establishment of irreversible meiotic exit.

## METHODS

### Growth conditions

For sporulation, strains were grown in synthetic complete medium (1xSC; 0.67% yeast nitrogen base without amino acids; 0.2% dropout mix with all amino acids; and, 2% glucose) at 30⁰C. Cells were then diluted 1:25 into synthetic complete acetate medium (1xSCA; 0.67% yeast nitrogen base without amino acids; 0.2% dropout mix with all amino acids; and, 2% acetate) at 30⁰C for 12-16 hrs.

### Strain construction

The *S. cerevisiae* strains used in this study are derivatives of W303 (*ade2-1 his3-11,15 leu2-3,112 trp1-1 ura3-1 can1-100*; Table S12). Genes were targeted for deletions and tagging using standard PCR-based lithium acetate transformation (Janke *et al.*, 2004). The *rim4-47A*, *rim4-27A*, and *rim4-20A* strains were originally obtained from the Berchowitz lab and the C-terminal mutations were transferred into our strain background by PCR amplification and transformation. The mutations were checked via sequencing. Genotypes of transformed strains were verified by PCR or by sequencing. Integrating plasmids *yomRuby2*-*TUB1* (yeast optimized mRuby2) were digested and integrated into the *URA3* locus (Markus *et al.*, 2015).

### Time-lapse imaging of budding yeast

For time-lapse imaging, 200µL of cells were concentrated and then adhered onto a concanavalin A-coated coverslip (Sigma; 1mg/mL in PBS) and inside a homemade chamber (Cairo *et al.*, 2022). To make a monolayer of cells, we spread the cells using a 5% agar plug (made with 1% potassium acetate) and then left the plug on the cells for 12 mins to allow time for the cells to adhere to the concanavalin A. 2 mls of the remaining pre-conditioned 1% potassium acetate was added dropwise to the chamber to float the agar plug. The plug was then removed and the chamber was put on the microscope immediately for imaging. All meiosis movies were initiated approximately 8 hours after the introduction of 1% potassium acetate, when the majority of cells were in prophase I.

Cells were imaged on a Nikon Ti2 microscope equipped with a Orca-fusion BT digital c-mos camera (Hamamatsu) and 60x or 100x oil-immersion objective lens. Images were acquired with GFP and Ruby filters with exposure time of 20-40ms and with neutral density filters transmitting 2-5% of light intensity. 5 z-stacks at 1.2µM z-stacks were acquired every 10 mins for 12-14 hrs. Brightfield images were acquired at 70ms at 5% neutral density. For image analysis, z-stacks were combined in a maximum intensity projection in NIS Elements software (Nikon). Fiji software (NIH) was used to create final images with adjustment of brightness and contrast.

### Statistical analysis

Statistical analysis was performed in Graphpad Prism. For meiotic timings, an unpaired, nonparametric Mann-Whitney test with computation of two-tailed exact P values was used. The two-sided Fisher’s exact test was used to analyze the percent of cells with additional SPBs.

### Mathematical Modeling

We numerically simulated a parsimonious mathematical model of the wiring diagram in Figure 3A, given by a system of nonlinear ordinary differential equations (Supplementary Information, Eqs. 3-14), using Python, corroborated by Mathematica’s NDSolve function employing a stiff solver (Wolfram Research, Inc., Mathematica, Version 13.3, Champaign, IL (2023)). Detailed description of the model, limitations and future work are given in the Supplementary Information and numerical values for all parameters and initial conditions are given in Tables S1-S10. The code used for the computational modeling described in this study is publicly available at https://github.com/abimarquez1211/Meiotic-Exit-Code.

## ACKNOWLEDGEMENTS

We thank the Lacefield Lab for careful reading of the manuscript. This work is supported by NSF award 2319006 to SL. Core facility support was provided by P20-GM113132 and 5P30CA023108 and the Genomics and Molecular Biology Shared Resource RRID:SCR021293.

## SUPPLEMENTARY INFORMATION

### Supplementary Figures

**Figures S1-S10** show the numerical solutions to all variables of the exit network, given by equations 3-14. In the main figure, only a subset of variables is shown for greater clarity.

**Figure S1** (related to Figure 3B). Dynamics for all exit network elements in wildtype cells. Concentrations are in arbitrary units (a.u.) on the vertical axes. The horizontal axes denote time since the start of meiosis II.

**Figure S2** (related to Figure 3D). Dynamics for all exit network elements in *clb1Δ* cells, with the concentration of Clb1 set to zero. Concentrations are in arbitrary units (a.u.) on the vertical axis. The horizontal axis denotes time since the start of meiosis II.

**Figure S3** (related to Figure 3E). Dynamics for all exit network elements in *clb4Δ* cells, with the concentration of Clb4 set to zero. Concentrations are in arbitrary units (a.u.) on the vertical axis. The horizontal axis denotes time since the start of meiosis II.

**Figure S4** (related to Figure 3F). Dynamics for all exit network elements in *clb3Δ* cells, with the concentration of Clb3 set to zero. Concentrations are in arbitrary units (a.u.) on the vertical axis. The horizontal axis denotes time since the start of meiosis II.

**Figure S5** (related to Figure 2F). Dynamics for all exit network elements in *clb1Δ clb4Δ* cells, with the concentration of Clb1 and Clb4 set to zero. Concentrations are in arbitrary units (a.u.) on the vertical axis. The horizontal axis denotes time since the start of meiosis II.

**Figure S6** (related to Figure 2F). Dynamics for all exit network elements in *clb1Δ clb3Δ* cells, with the concentration of Clb1 and Clb3 set to zero. Concentrations are in arbitrary units (a.u.) on the vertical axis. The horizontal axis denotes time since the start of meiosis II.

**Figure S7** (related to Figure 2F). Dynamics for all exit network elements in *clb3Δ clb4Δ* cells, with the concentration of Clb3 and Clb4 set to zero. Concentrations are in arbitrary units (a.u.) on the vertical axis. The horizontal axis denotes time since the start of meiosis II.

**Figure S8** (related to Figure 5A). Dynamics for all exit network elements in *ama1Δ* cells, with the concentration of Ama1 set to zero. Concentrations are in arbitrary units (a.u.) on the vertical axis. The horizontal axis denotes time since the start of meiosis II.

**Figure S9** (related to Figure 4B). Dynamics for all exit network elements in *rim4-47AΔ* cells, with delayed Rim4 degradation. Concentrations are in arbitrary units (a.u.) on the vertical axis. The horizontal axis denotes time since the start of meiosis II.

**Figure S10.** Dynamics for all exit network elements in cells that lack autophagy (no Rim4 degradation). Concentrations are in arbitrary units (a.u.) on the vertical axis. The horizontal axis denotes time since the start of meiosis II.

**Figure S11.** Sigmoidal dynamics of mRNA-Rim4 dissociation and Rim4 clearance for WT and representative Rim4 mutants, using the phenomenological description adopted in this work.

**Figure S12.** Total CDK-Clb concentration and threshold crossing for different Rim4 mutants, recapitulating *rim4-20A, rim4-27A,* and *rim4-47A* experimental results for the duration of metaphase II.

### 1 Supplementary Information: Mathematical Model for Meiotic Exit Network

#### 1.1 Model Description

##### 1.1.1 Minimal Network Model

The mathematical model given by the system of nonlinear ordinary differential equations, Eqs. 3-14, describes the wiring diagram in Fig. 3A, as a minimal model of exit from meiosis II established in this work. This model builds on that of Okaz *et al.* [38] which described the irreversible transition from prophase I to metaphase I.

The rate of change of the concentration of each variable with time is described by a nonlinear, ordinary differential equation (ODE) given by the relevant biochemical reaction kinetics, described below. To numerically simulate the equations, we used Python^1^, corroborated by Mathematica’s `NDSolve` function employing a stiff solver (Wolfram Research, Inc., Mathematica, Version 13.3, Champaign, IL (2023)). Values for shared parameters were based on Okaz *et al.* [38], while values for new parameters were determined in a trial and error process aimed at reproducing the behavior of wild-type and mutants studied in this work, consistent with other works [13, 12, 41, 37, 50, 26, 36, 16]. Rate constants are in units of min^-1^; concentrations are in arbitrary units (a.u.). Parameter values and initial conditions are reported in Tables S1-10.

The key components of this network are cyclin-dependent kinase (Cdk1) heterodimers, Cdk1-Clb1, Cdk1-Clb3 and Cdk1-Clb4; polo kinase, Cdc5 (with role in Cdc14 release from and re-sequestration to the nucleolus, not accounted for in mathematical model); Ndt80, which transcribes Clb1, Clb3, Clb4, Cdc5, Cdc20, and itself through a positive feedback loop; ubiquitin ligase APC/C (anaphase promoting complex), where APC/C alone is unable to target cyclins for degradation, rather its activation is dependent on the binding of cofactors; Ama1 and Cdc20, activators of APC; Rim4, translational repressor of Ama1 and Clb3 mRNA, which is degraded by autophagy in meiosis II (after being phosphophorylated by Ime2, not accounted for in our mathematical model).

We explicitly take into account APC/C activation by Cdc20, following [14, 41] (Eq. 9, see also below). However, in the case of APC/C^Ama1^, as in [38], we assume that the concentration of APC core is not rate-limiting, and therefore the concentration of APC/C^Ama1^ follows that of Ama1 in our model (Eq. 13). Similarly, it is assumed that Cdk1 subunits are present in excess, and their concentration is not rate-limiting; hence the concentrations of Cdk1-Clb1, Cdk1-Clb3, and Cdk1-Clb4 follow that of the cyclin subunits (Eqs. 3, 4, 5). Synthesis and degradation reactions are approximated by mass action kinetics. Reactions describing activation and inactivation by phosphorylation are described according to zeroth order Goldbeter-Koshland kinetics (Eqs. 8, 9, and Eqs. 12, 13), following previous works [25, 38]. Cdk1-Clb1/Clb4/Clb3 inhibit APC/C^Ama1^ by multisite phosphorylation (Eqs. 12). The most active form of Ama1 is unphosphorylated. Phosphorylation reduces but does not completely suppress APC/C^Ama1^ activity. Therefore, Ama1-dependent proteolysis (Eqs. 3, 4, 5, 10, 11) is proportional to the concentration of active, unphosphorylated Ama1 ([Ama1]) as well as to the total concentration of Ama1 ([Ama1_T_] = [Ama1-P]+[Ama1]).

Rim4 is phosphorylated by Ime2, leading to its disassembly from amyloid-like aggregates, release of bound mRNAs, and targeting by autophagy for degradation in metaphase II. Importantly, for the network modeled here, this allows Ama1 mRNA which is sequestered by Rim4 at the start of meiosis II to be translated, giving rise to a surge of Ama1 protein (Eq. 13). APC/C^Ama1^ is held inactive due to Cdk1-Clb1/Clb4 inhibition by phosphorylation (Eq. 12). As Cdk1-Clb1/Clb4 levels rise in metaphase II, they phosphorylate and activate APC/C^Cdc20^ (Eqs. 8, 9). Active APC/C^Cdc20^ drives the cells into anaphase II by targeting proteins like Clb1/4 for degradation, decreasing Cdk1-Clb1/Clb4 activity (Eqs. 3, 4, 5). With decreased Cdk1-Clb1/Clb4, APC/C^Ama1^ becomes active and degrades Ndt80, Cdc5, and more Clb1/Clb4 and Clb3 (Eqs. 3-6 10-11). Without these regulators, the cell exits meiosis. Clb3, also expressed with release of Clb3 mRNA from Rim4-mRNA aggregates (Eq. 7), extends the duration of APC/C^Cdc20^ activity, but otherwise does not significantly affect the timing of exit for the chosen numerical values of parameters governing Clb3 kinetics in our current model (motivated by experimental results; see main text). Also, in the present model, Cdk1-Clb4 (and later Cdk1-Clb3), in addition to Cdk1-Clb1, inhibits APC/C^Ama1^ activity by phosphorylation (Eq. 12). As such, the role of the unknown inhibitor of Ama1 in Okaz *et al.* [38] is played by Clb4.

It has been established that the negative feedback loop between Cdk1-Clb (here, given by Clb1/Clb4/3) and APC/C^Cdc20^ gives rise to oscillatory dynamics [37, 50, 33, 2]. However, oscillations require an intermediate step, which serves to provide a delay between Cdk1-Clb1/4 activation of APC/C^Cdc20^ and APC/C^Cdc20^ inhibition (via ubiquitination) of Cdk1-Clb1/4. In our model, this intermediate step is achieved through Cdk1-Clb1/4 phosphorylation of the APC/C core (Eq. 8), followed by its activation through binding to Cdc20 (Eq. 9) [41]. Additionally, we allow for APC/C^Cdc20^ degradation of Cdc20 (via self-ubiquitination) (Eq. 7). Further mechanistic details of this feedback loop, which include the inhibitory role of phosophorylation of Cdc20 protein [15, 31], could also be important and are the subjects of future work.

##### 1.1.2 Cdk1-Clb threshold

With inhibition of autophagy, it has been shown that several rounds of spindle formation, spindle elongation, and chromosome segregation ensue [52]. This is recapitulated as damped oscillations in Cdk1-Clb concentrations in the exit network dynamics. Indeed, in the network model considered here, the Cdk1-Clbs play redundant roles, motivating our consideration of the *total* Cdk1-Clb concentration in the network dynamics. Since the decline in Cdk1-Clb activity is critical for exiting meiosis, we consider the time it takes from the start of meiosis II (t = 0) for the total Cdk1-Clb concentration level to fall below a threshold, given by approximately one third of the peak total Cdk1-Clb level in WT. This time interval is compared with the experimental results on the duration of metaphase II in WT and mutant strains (Table S11). Although this threshold has been determined by trial and error, we emphasize that a single threshold value yields results that are consistent with experiments for WT and seven mutant phenotypes.

##### 1.1.3 mRNA-Rim4 dynamics and Rim4 clearance

We emphasize that a quantitative description of the spatiotemporally coordinated mechanisms of mRNA sequestration through complex formation with Rim4 amyloid aggregates, followed by Rim4 phosphorylation-mediated release of mRNA and degradation of Rim4 via authophagy [52, 28] is beyond the scope of the current work. Rather, here, we describe the dissociation of Rim4-mRNA complexes and Rim4 clearance phenomenologically, given by a single, sigmoidal function of time, 0≤ *f(t; T, τ_p_*) ≤ 1, where *f(t; T, τ_p_*) = 1 corresponds to no protein translation (i.e., mRNA is fully sequestered in Rim4 aggregates) and no Rim4 degradation (no autophagy). Specifically, we use the following functional form:

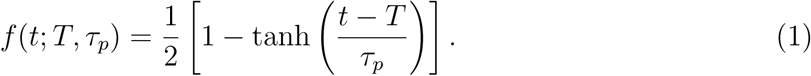

Cooperative models [34, 30, 8, 44, 32, 18, 20, 19, 40] of multisite phosphorylation kinetics applied to Rim4 amyloid aggregates could plausibly describe the transition from the mRNA-bound (and unphosphorylated) to the mRNA-unbound (and phosphorylated) state. These models display similar sigmoidal functional behavior as the phosphorylation level of Rim4 increases with time, motivating our description.

In Fig. S11, we show the role of the parameter *T*, which determines the time of onset of Rim4-mRNA dissociation (and Rim4 clearance), and the parameter τ_p_, which determines the time scale of Rim4-mRNA dissociation, both of which in principle depend on the details of the cooperative, multisite phosphorylation kinetics. In this figure, parameters given by (T = 20 min, 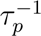= 0.2 min^-1^), corresponds to WT (solid gray), while for parameters given by (*T* = 43 min, 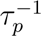 = 0.2 min^-1^) mRNA-Rim4 dissociation is delayed with respect to WT (dashed gray), and with (*T* = 43 min, 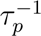 = 0.075 min^-1^) there is both a delay and slower dissociation. A mechanistic description of this process – including the balance between kinase and phosphatase activities of Ime2 and Cdc14, respectively, in determining the overall phosphorylation level of Rim4 – is the subject of future experimental and modeling studies. Finally, as shown in Fig. 4A, we assume that for Rim4 mutants resulting in its reduced phosophorylation, dissociation of mRNA-Rim4 complex and Rim4 clearance is delayed (T increases) while the rate of dissociation/clearance (1/τ_*p*_) is approximately unchanged. These assumptions are plausibly achieved within mechanistic models of cooperativity, which we plan to further explore in future work. In Figure S12 we show the numerical solution for the total Cdk1-Clb level as a function of time for different values of T, demonstrating that as mRNA-Rim4 dissociation is delayed, the threshold crossing occurs later in meiosis II, consistent with the experimental observations of prolonged metaphase II (Fig. 4C), and eventual transition to oscillatory dynamics.

We note that degradation of Rim4 is relevant for the dynamics of the exit network only in so far as ensuring that released mRNA is not re-sequestered. Even if mRNA does not rebind to phosphorylated Rim4, with release of Cdc14 from the nucleolus during anaphase II, its phosphatase activity could result in sufficient dephosphorylation of Rim4 for mRNA rebinding to occur. The present model assumes no rebinding of mRNA (equivalent to rapid clearance of Rim4). Separating the effects of mRNA rebinding and Rim4 clearance via autophagy on the dynamics of exit is the subject of future work.

The rate at which Clb3 and Ama1 protein are produced from mRNA sequestered by Rim4 is given in terms of Eq. 1 as

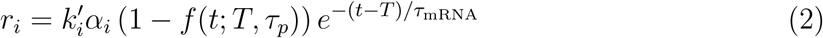

where i = Clb3 or Ama1. The parameters (α_Clb3_, α_Ama1_) reflect the amount of sequestered mRNA, and (*k*^’^_CIb1_*k*^’^_Ama1_) are the translation rates. The exponentially decaying function accounts for the finite lifetime of released mRNA. The mRNA lifetime is taken to be τ_mRNA_ ≈ 25 minutes (= average mRNA lifetime in yeast, from [21]).

#### 1.2 Limitations and Future Work

Regulation of mitosis in budding yeast has been the subject of extensive experimental studies and related theoretical modeling, where input of new experimental data has resulted in steady refinement of mathematical models of the underlying cell cycle circuitry [49, 13, 12]. In contrast, mathematical modeling of the regulation of meiosis is less well-characterized [15, 38, 6]. In this work, we have presented a parsimonious model of exit from meiosis II that captures experimental observations in budding yeast ranging from wild-type to single and double mutations with a single set of parameter values. We similarly expect further refinements of our model, informed by future experiments.

Key limitations of our current model and possible extensions are discussed below:

1. As described above, we have not attempted to describe the mRNA-Rim4 dynamics mechanistically. Rather, we have addressed the role of Rim4 in regulating exit – through the delay or suppression of release of mRNA – phenomenologically. This component of the network dynamics can be further expanded in the following ways:

i. Cooperative models of multisite phosphorylation governing mRNA-Rim4 association and dissociation would provide a principled description, that can be related to RNA-binding properties of Rim4 and its mutants [34, 30, 8, 44, 32].
ii. RNA/protein (RNP) bodies are an important class of membrane-less organelles that bind and regulate RNA, thereby ensuring the proper execution of a variety of cellular regulatory processes in a spatially and temporally controlled manner [10, 58, 45, 42, 9]. Increasing evidence suggests that the assembly of RNP bodies can be described in terms of liquid-liquid phase transitions; furthermore, intrinsically disordered protein motifs are commonly found in RNP bodies [53, 51, 56]. Therefore, should future experiments reveal liquid-liquid phase separation as the mechanism governing the interaction of Rim4 and its mRNA binding partners, the dynamics of phase separation and its role in regulating exit will need to be treated in the theoretical model [55].
2. The inherent stochasticity arising from thermal energy of biomolecules gives rise to intrinsic fluctuations in the concentrations of components and rates of reactions comprising biological circuits, thereby constraining the accuracy of biochemical processes [47, 17, 7, 35, 43, 48, 39]. A major source of intrinsic noise can be attributed to low copy numbers of mRNA molecules in gene-protein regulatory networks [35]. Here, we have adopted a deterministic model and numerical simulation approach to describe the regulatory network in Fig. 3A. Future stochastic simulations will address fluctuations in protein and mRNA numbers [23, 24, 4, 3, 1, 11, 22], allowing comparison of numerical results with experimentally measured variabilities in cell fates and timing of meiotic exit. This will be especially relevant for comparison with our experimentally measured population-level distributions of the timing exit and cell fates in meiosis II, where the decline and threshold crossing of noisy Cdk1-Clbs level will give rise to variability in outcomes for exit. Related to this, recent work has suggested that fluctuations in protein numbers are mitigated by phase-separated compartments, thereby reducing noise and enhancing the robustness of biological systems [46, 29, 57], suggesting a further role for mRNA-Rim4 aggregates.
3. Refinements to the network connectivity that are not treated in the present work include other kinase and phosphatase activities (notably Ime2 and Cdc14, respectively), with possible impact on the biochemical oscillator module (presently given by Cdk1Clbs and APC/C^Cdc20^) as well as dynamics of mRNA-Rim4 dissociation mediated by the phosphorylation state of Rim4. Additionally, Clb3 does not appear to play a central role in the dynamics of meiotic exit, despite the fact that its mRNA is a target of Rim4. This surprising experimental finding, reproduced in the mathematical model, may point to other role(s) for Clb3 in meiosis II that are not considered here.
4. Quantitative computational models in systems biology typically involve many free parameters, and directly measuring in vivo or in vitro biochemical parameters is challenging. The difficulty in assigning values of these parameters hinders model development and theoretical analysis. Earlier work has shown that the quantitative behavior of many different classes of nonlinear multiparameter models is more sensitive to changes in certain combinations of parameters than others (“stiff” versus “sloppy” parameter subspaces) [27]. Similar analyses on models of the budding yeast cell cycle [5, 54] have revealed an effective reduction in the number of degrees of freedom of the models, and could provide further insight into refinements of the model of meiotic exit presented here.

#### 1.3 Model Equations

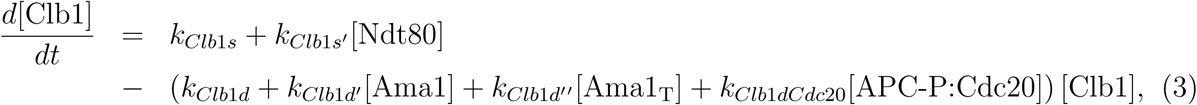

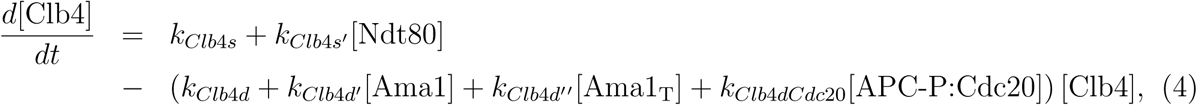

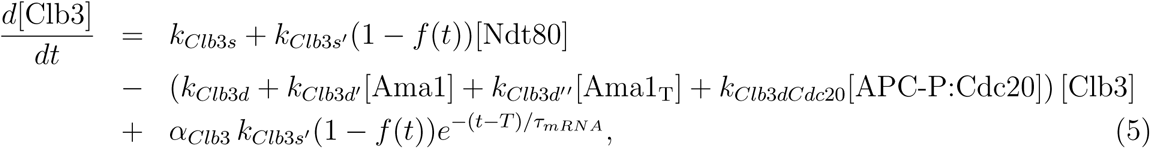

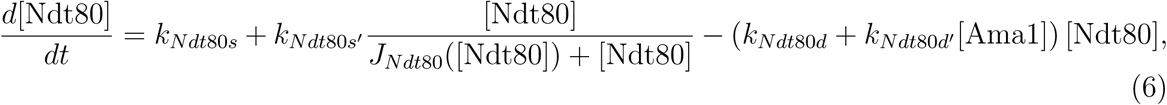

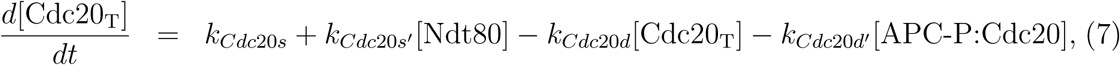

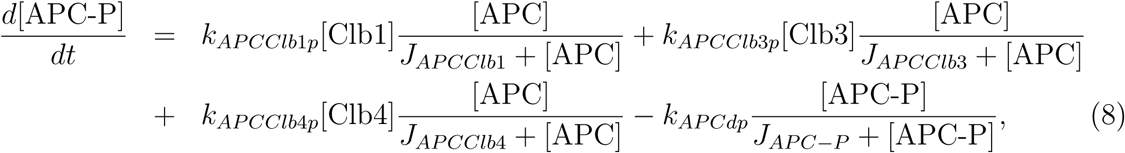

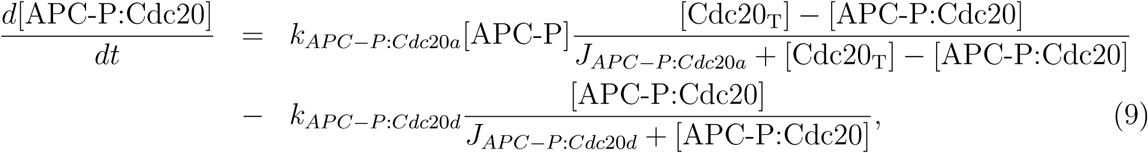

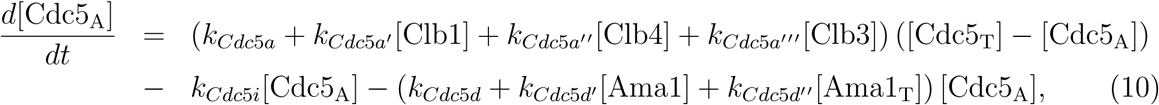

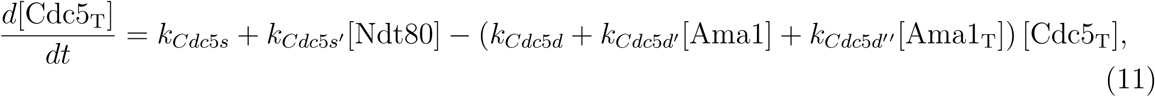

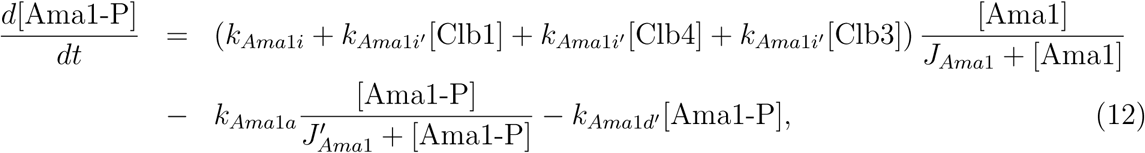

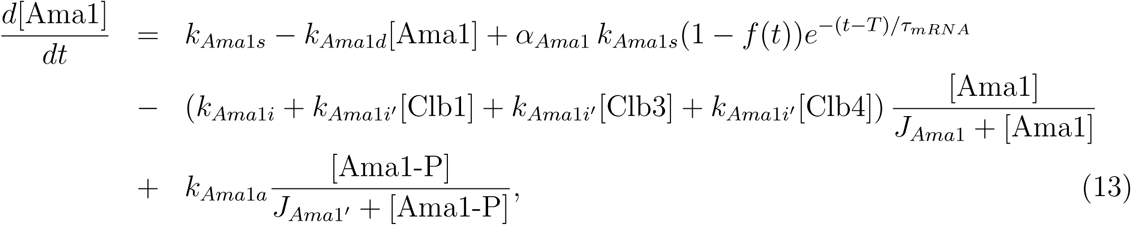

Additionally,

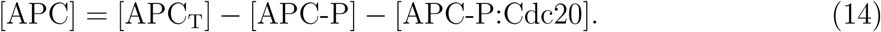

#### 1.4 Model Parameters, Initial Conditions

**Table S1:** Clb1 Parameters.

| Parameter | Value |
| --- | --- |
| $k_{Clb1s}$ | 0.04 |
| $k_{Clb1s'}$ | 0.024 |
| $k_{Clb1d}$ | 0.1 |
| $k_{Clb1d'}$ | 0.5 |
| $k_{Clb1d''}$ | 0.02 |
| $k_{Clb1dCdc20}$ | 0.8 |

**Table S2:**
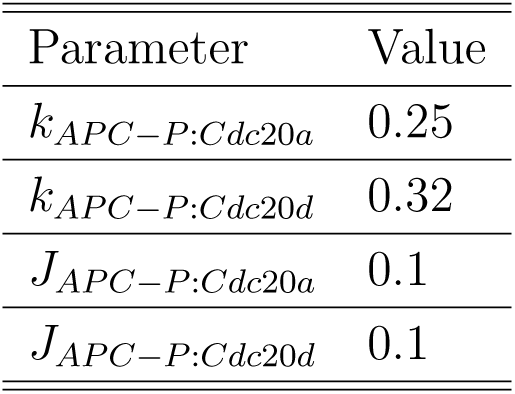
APC/C^Cdc20^ Parameters.

**Table S3:** Clb3 Parameters.

| Parameter | Value |
| --- | --- |
| $k_{Clb3s}$ | 0.002 |
| $k_{Clb3s'}$ | 0.000114 |
| $k_{Clb3d}$ | 0.34 |
| $k_{Clb3d'}$ | 0.2 |
| $k_{Clb3d''}$ | 0.02 |
| $k_{Clb3dCdc20}$ | 0.4 |
| $\alpha_{Clb3}k_{Clb3s'}$ | 0.4 |
| $\tau_{mRNA}$ | 25 |

**Table S4:** Cdc5 Parameters.

| Parameter | Value |
| --- | --- |
| $k_{Cdc5a}$ | 0.1 |
| $k_{Cdc5i}$ | 2 |
| $k_{Cdc5d}$ | 0.04 |
| $k_{Cdc5d'}$ | 0.003 |
| $k_{Cdc5d''}$ | 0.002 |
| $k_{Cdc5a'}$ | 1.2 |
| $k_{Cdc5a''}$ | 1.2 |
| $k_{Cdc5a'''}$ | 2 |
| $k_{Cdc5s}$ | 0.004 |
| $k_{Cdc5s'}$ | 0.027 |

**Table S5:** Clb4 Parameters.

| Parameter | Value |
| --- | --- |
| $k_{Clb4s}$ | 0.0008 |
| $k_{Clb4s'}$ | 0.009 |
| $k_{Clb4d}$ | 0.02 |
| $k_{Clb4d'}$ | 0.2 |
| $k_{Clb4d''}$ | 0.02 |
| $k_{Clb4dCdc20}$ | 0.2 |

**Table S6:** Ama1 Parameters.

| Parameter | Value |
| --- | --- |
| $k_{Ama1i}$ | 0.005 |
| $k_{Ama1i'}$ | 0.5 |
| $J_{Ama1}$ | 0.1 |
| $k_{Ama1a}$ | 0.1 |
| $J_{Ama1'}$ | 0.1 |
| $k_{Ama1s}$ | 0.04 |
| $k_{Ama1d}$ | 0.02 |
| $k_{Ama1d'}$ | 0.02 |
| $\alpha_{Ama1}k_{Ama1s}$ | 0.4 |
| $\tau_{mRNA}$ | 25 |

**Table S7:** Ndt80 Parameters.

| Parameter | Value |
| --- | --- |
| $k_{Ndt80s}$ | 0.01 |
| $k_{Ndt80s'}$ | 2 |
| $k_{Ndt80d}$ | 0.093 |
| $k_{Ndt80d'}$ | 0.013 |
| $J_{Ndt80}(x)$ | $a + b/x$ |
| $(a, b)$ | $(8.5, 2.5 \times 10^4)$ |

**Table S8:**
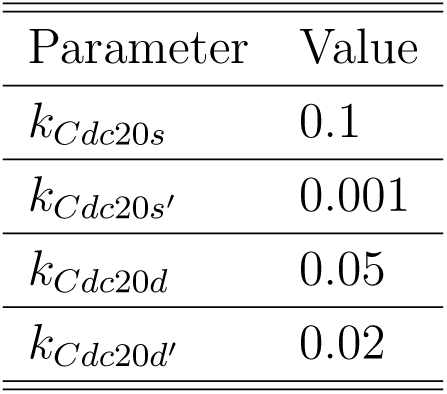
Cdc20 Parameters.

| Parameter | Value |
| --- | --- |
| $k_{Cdc20s}$ | 0.1 |
| $k_{Cdc20s'}$ | 0.001 |
| $k_{Cdc20d}$ | 0.05 |
| $k_{Cdc20d'}$ | 0.02 |

**Table S9:** APC/C Parameters.

|  |  |
| --- | --- |
| $k_{APCClb1p}$ | 0.09 |
| $k_{APCClb4p}$ | 0.09 |
| $k_{APCClb3p}$ | 0.09 |
| $J_{APCClb1}$ | 0.001 |
| $J_{APCClb4}$ | 0.001 |
| $J_{APCClb3}$ | 0.001 |
| $k_{APCdp}$ | 0.072 |
| $J_{APC-P}$ | 0.01 |
| $[APC_T]$ | 5 |

**Table S10:** Initial Conditions.

|  |  |
| --- | --- |
| Clb1 | 1.125 |
| Clb4 | 0.12 |
| Clb3 | 0 |
| Ndt80 | 5 |
| Cdc20<sub>T</sub> | 2 |
| APC-P | 0 |
| APC-P:Cdc20 | 0.1 |
| Cdc5<sub>A</sub> | 0.25 |
| Cdc5<sub>T</sub> | 0.75 |
| Ama1-P | 0 |
| Ama1 | 0 |

**Table S11:** Timing of Cdk1-Clb threshold crossing, t_threshold_ (model), and duration of metaphase II, t_metaphase II_ (experiment)

| Strain | $t_{\text{threshold}}$ (min) | $t_{\text{metaphase II}}$ (min) |
| --- | --- | --- |
| WT | 34 | $38 \pm 18$ |
| *clb1*Δ | 48 | $52 \pm 33$ |
| *clb4*Δ | 40 | $37 \pm 13$ |
| *clb3*Δ | 27 | $36 \pm 16$ |
| *ama1*Δ | 50 | |
| *rim4-47A-1* ($T=38$) | 56 | $56 \pm 37$ |
| *rim4-47A-2* ($T=43$) | 75* | - |
| *rim4-27A* | 48 | $47 \pm 15$ |
| *rim4-20A* | 40 | $41 \pm 12$ |

\* onset of oscillations

#### 1.5 Supplementary Figures

For completeness, Figures S1-S10 show the numerical solutions to all variables of the exit network, given by Eqs. (3)-(14), using initial conditions given in Table S10 for wild-type. For each deletion, if the initial concentration is nonzero, it is set equal to zero. Concentrations are in arbitrary units (a.u.). The horizontal axis denotes time since the start of meiosis II. In the main manuscript, only a subset of the solutions are shown for greater clarity. Figure S11 shows the mRNA-Rim4 dissociation dynamics for different timings of mRNA release, and Figure S12 shows the resulting total Cdk1-Clb concentration and threshold crossing.

**Figure S1:**
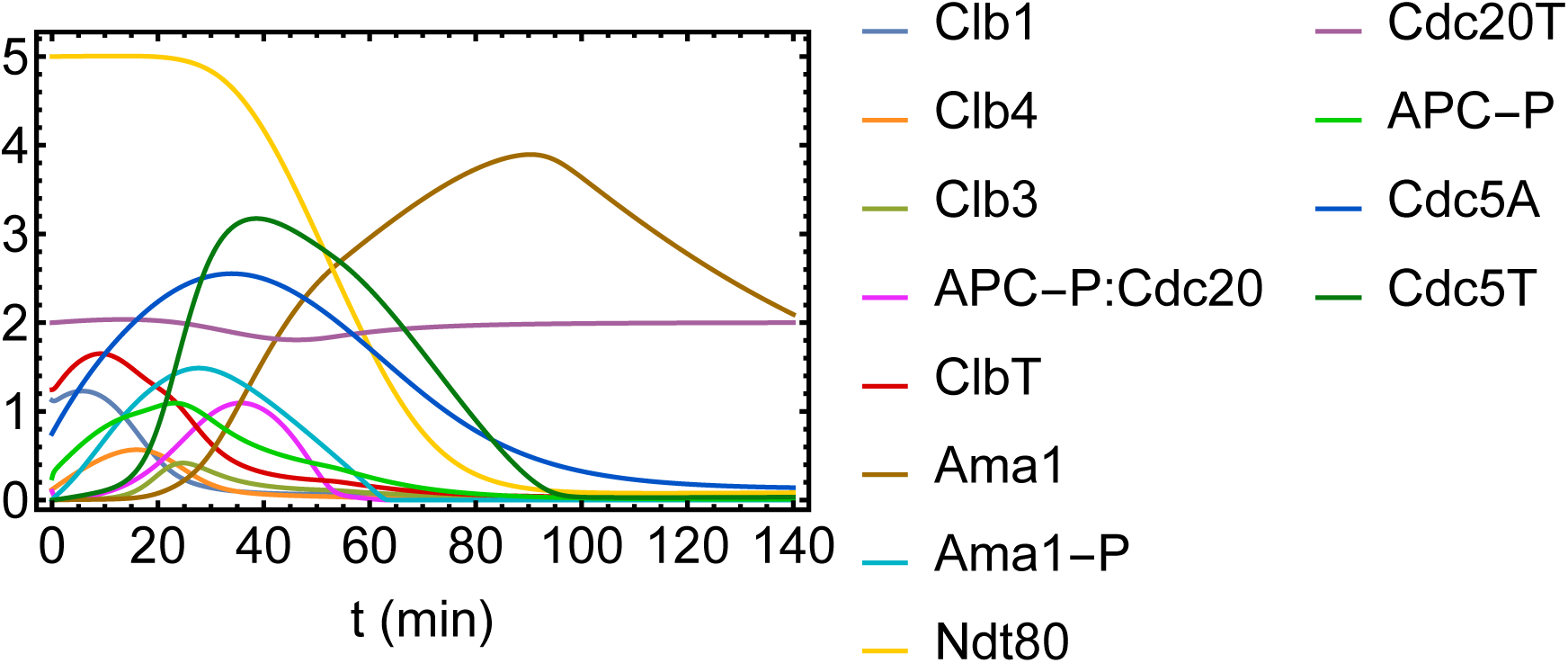
Wild-type.

**Figure S2:**
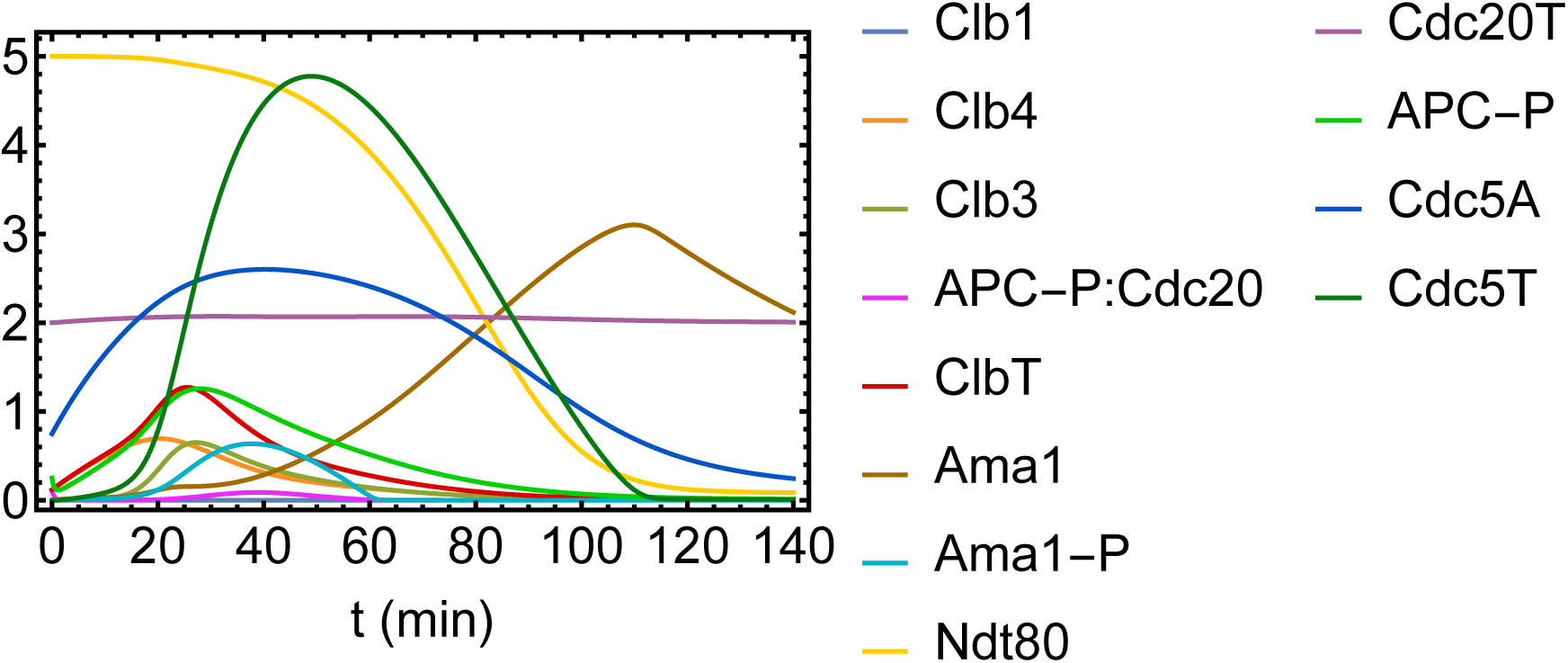
Clb1 deletion.

**Figure S3:**
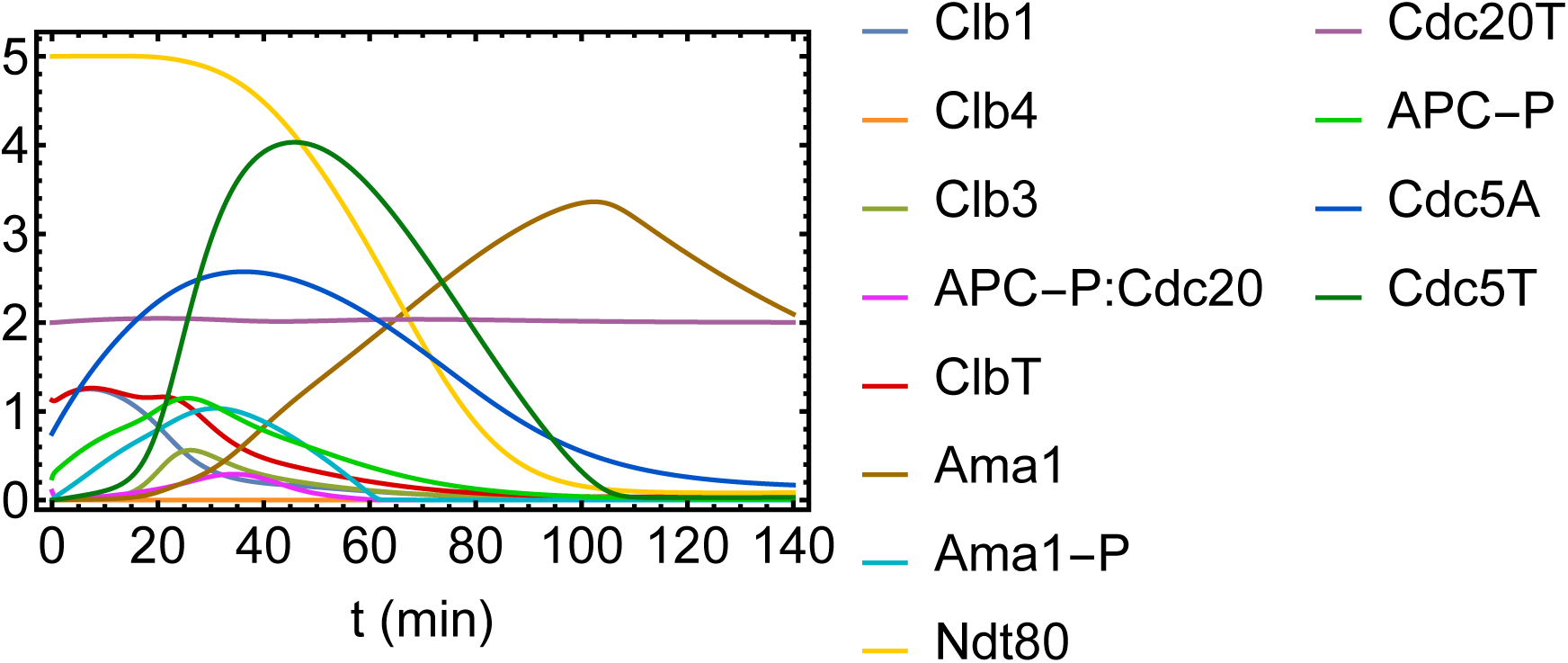
Clb4 deletion.

**Figure S4:**
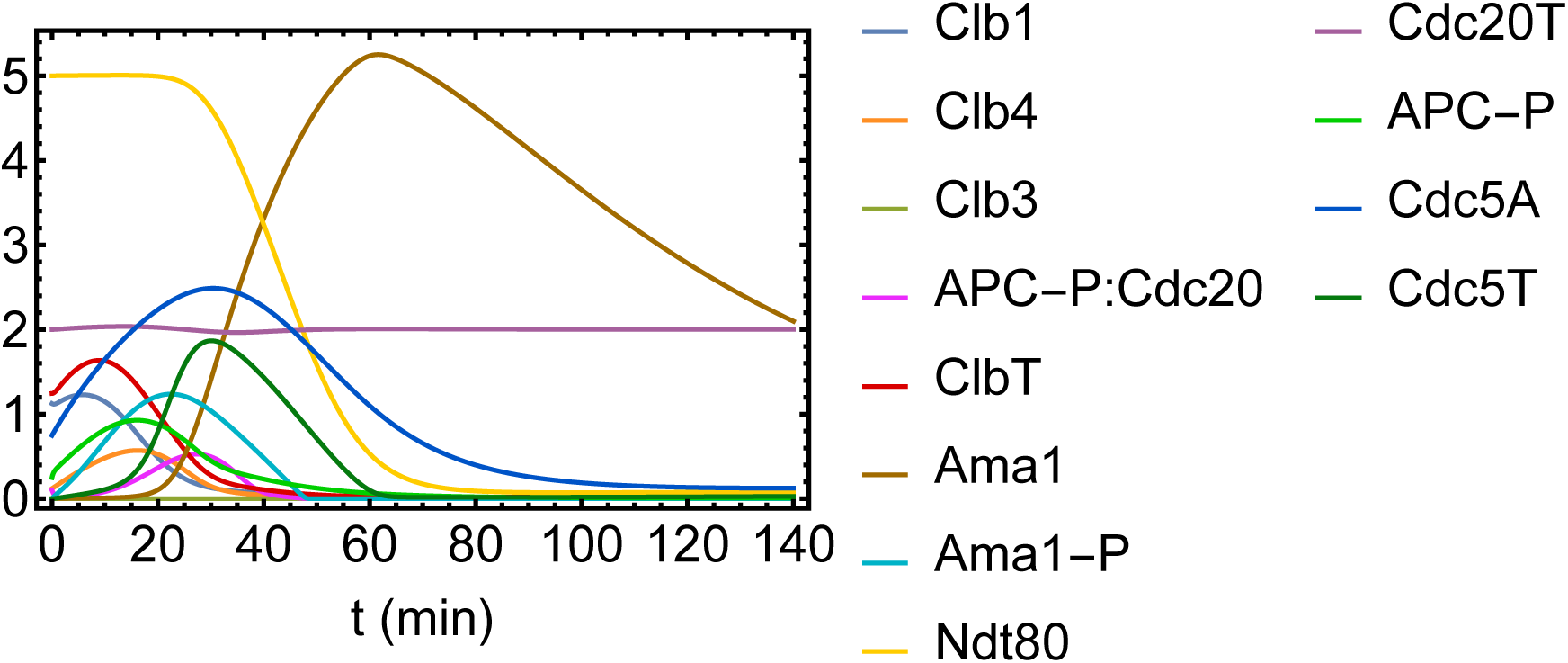
Clb3 deletion.

**Figure S5:**
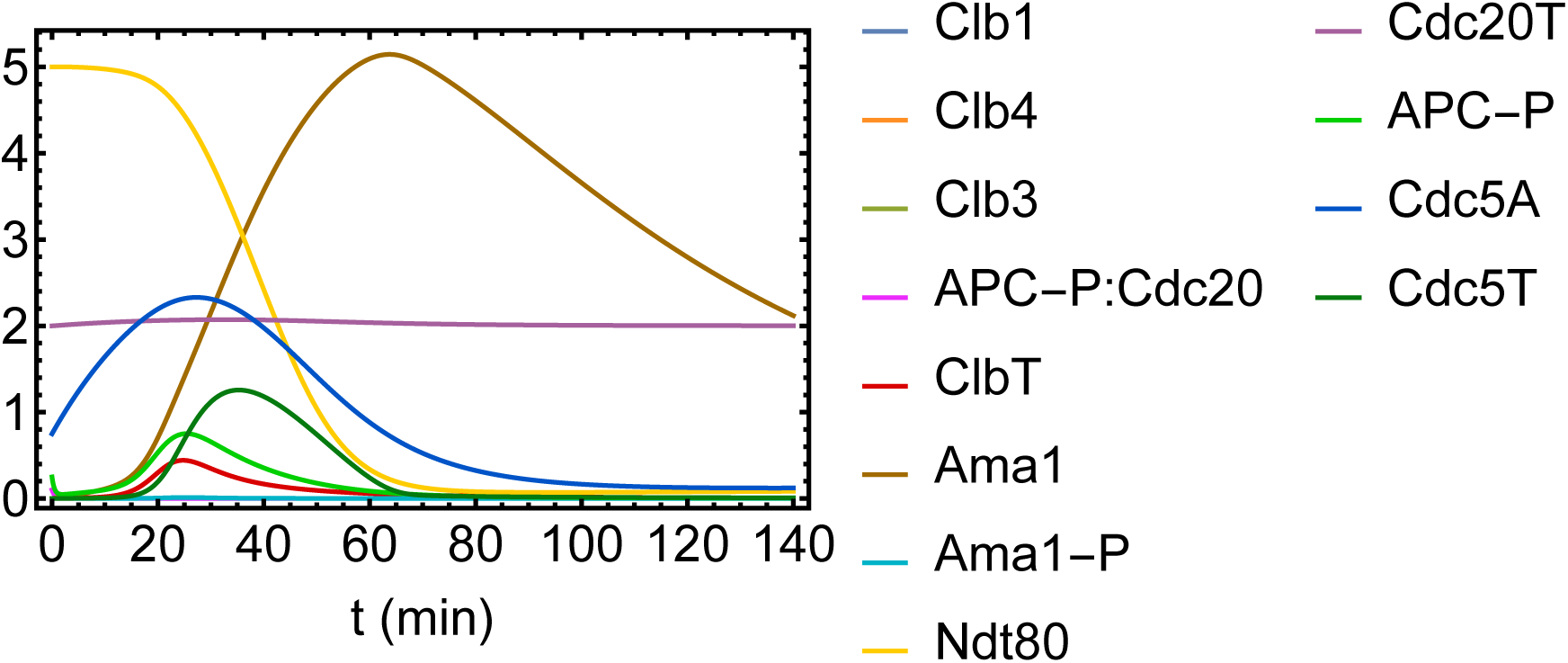
Clb1-Clb4 deletion.

**Figure S6:**
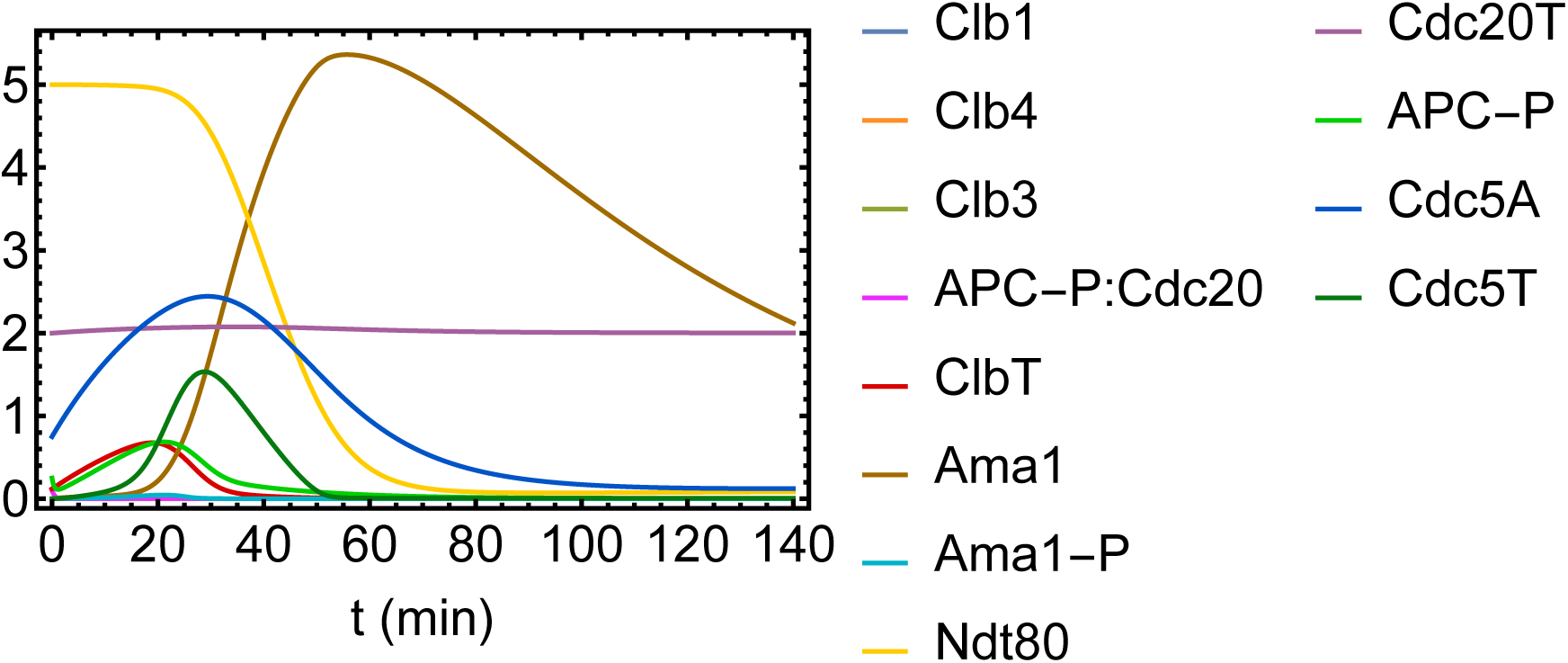
Clb1-Clb3 deletion.

**Figure S7:**
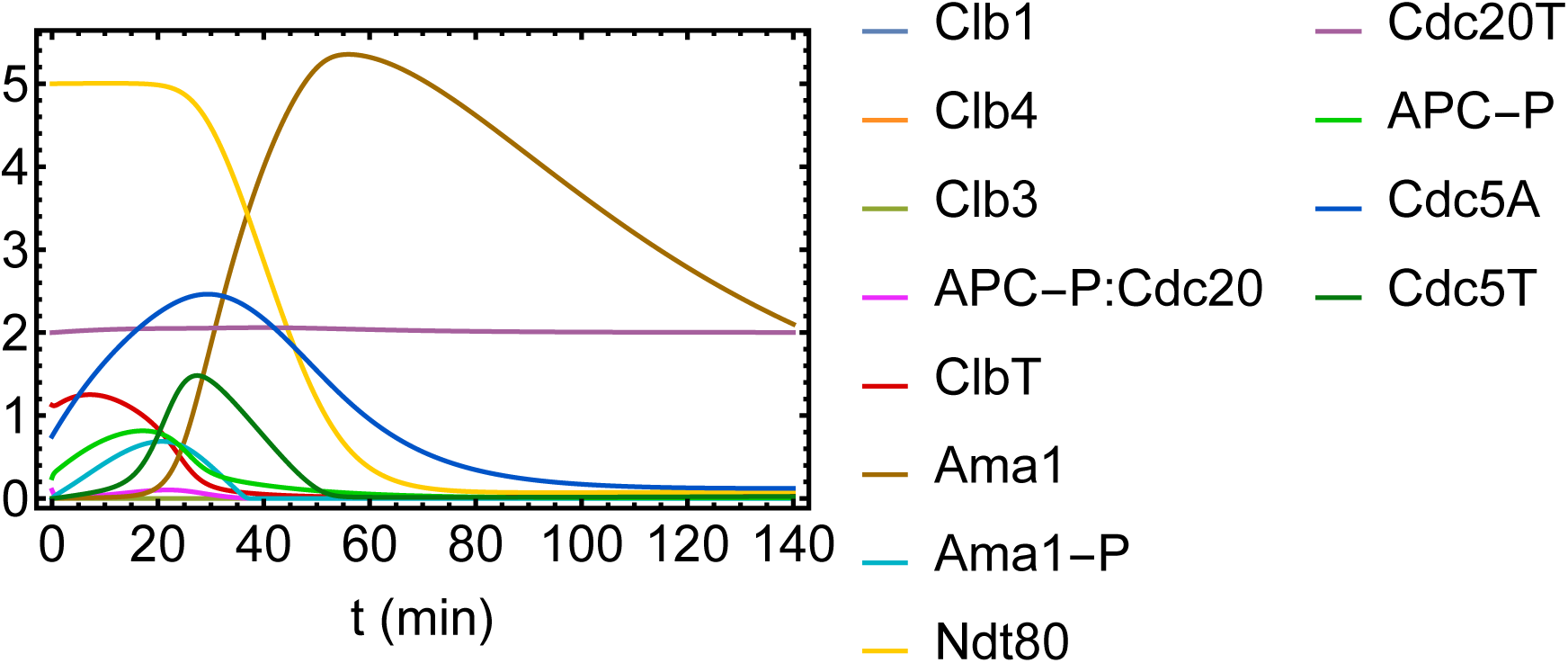
Clb3-Clb4 deletion.

**Figure S8:**
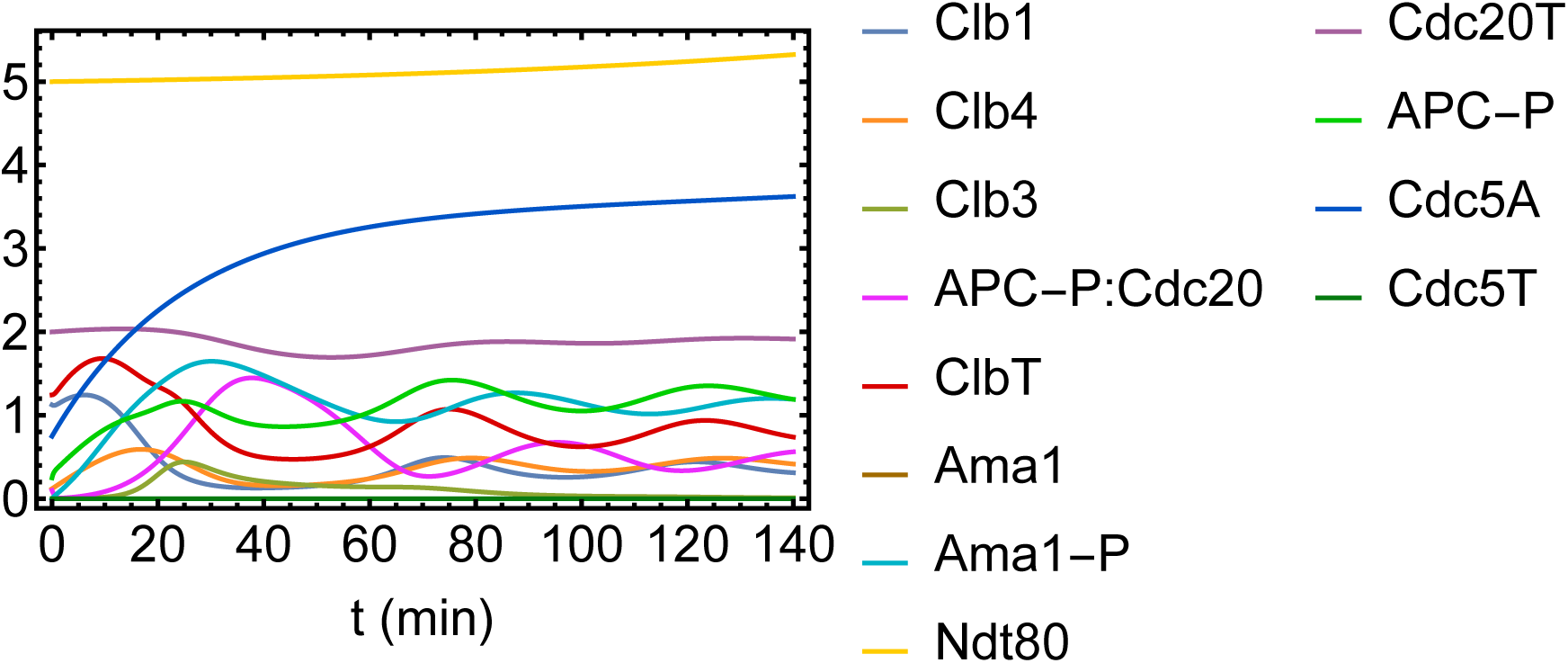
Ama1 deletion.

**Figure S9:**
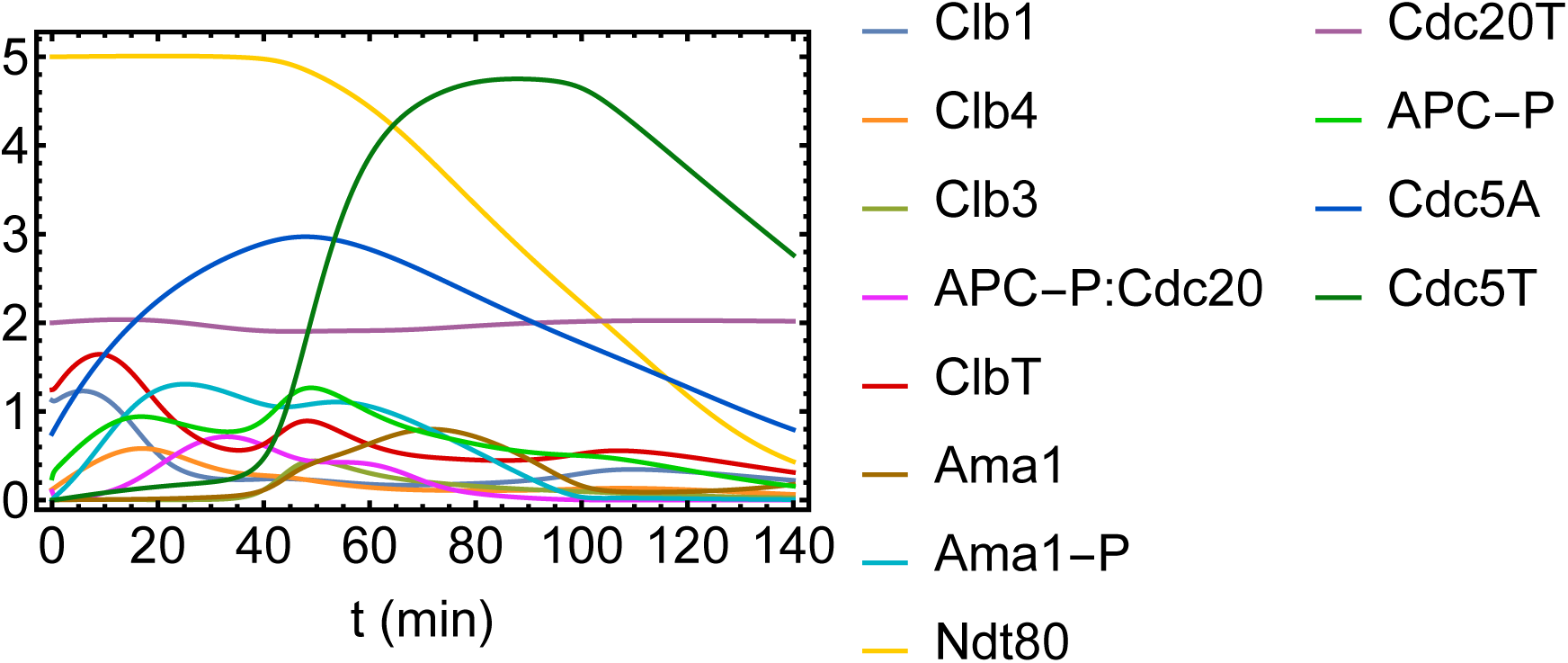
*rim4-47A-2* mutant.

**Figure S10:**
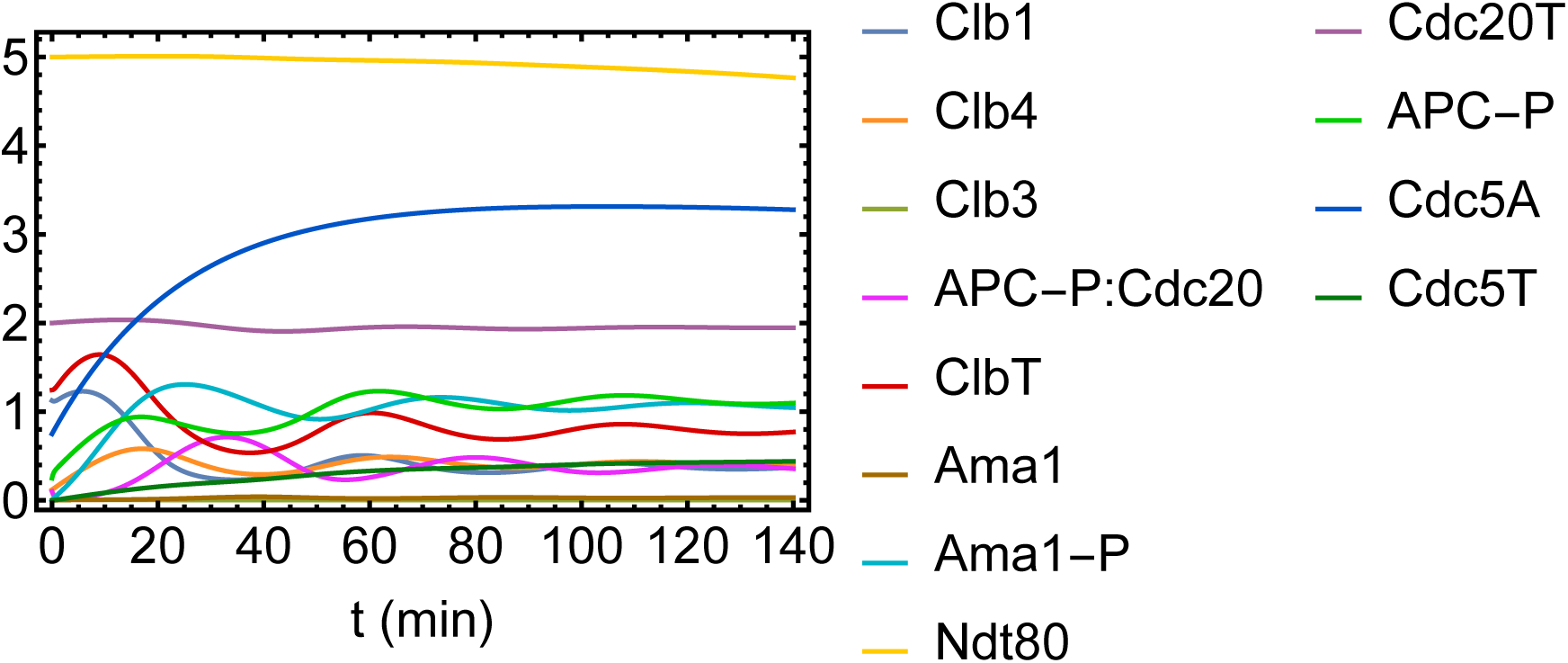
No autophagy (*f(t; T, τ_p_*) = 1)

**Figure S11:**
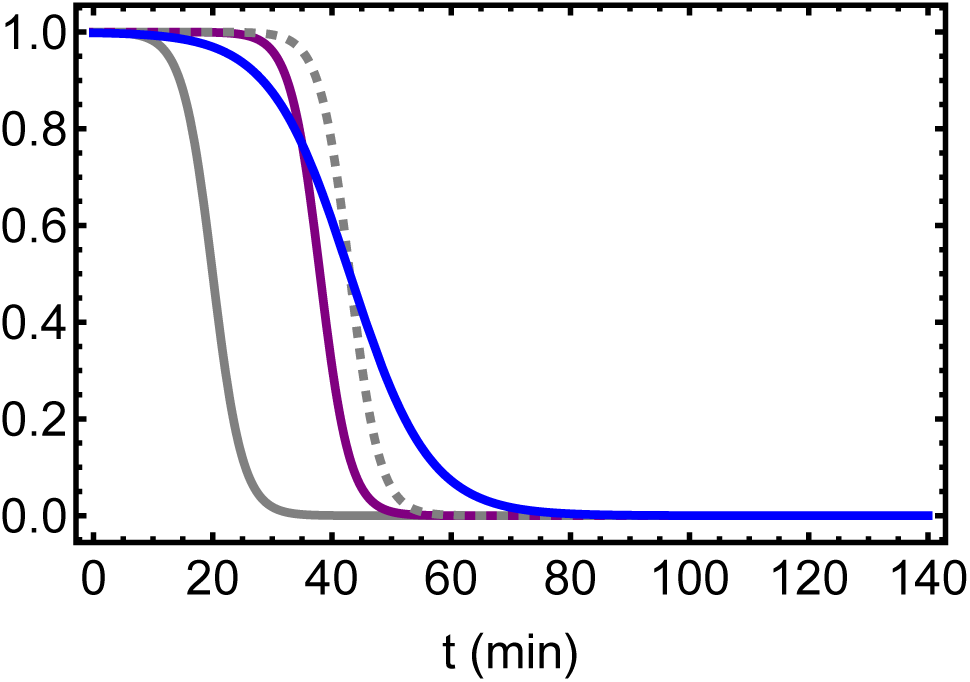
mRNA-Rim4 dissociation and Rim4 clearance given by Eq. 1. Solid gray: T = 20 min, 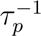 = 0.2 min^-1^ (WT). Solid purple: T = 38 min, 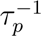 = 0.2 min^-1^. (*rim4-47A-1*). Dashed gray: T = 43 min, 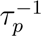 = 0.2 min^-1^. Solid blue: T = 43 min, 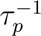 = 0.075 min^-1^ (*rim4-47A-2*). Solid blue: T = 43 min, 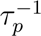 = 0.075 min^-1^. To describe Rim4 phosphorylation mutants, we assume 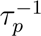(= 0.2 min^-1^) is fixed, and T increases with respect to WT. The horizontal axis is time in meiosis II, and on the vertical axis, a value of 1.0 corresponds to complete sequestration of mRNA by Rim4, and 0.0 corresponds to complete release of mRNA and Rim4 clearance.

**Figure S12:**
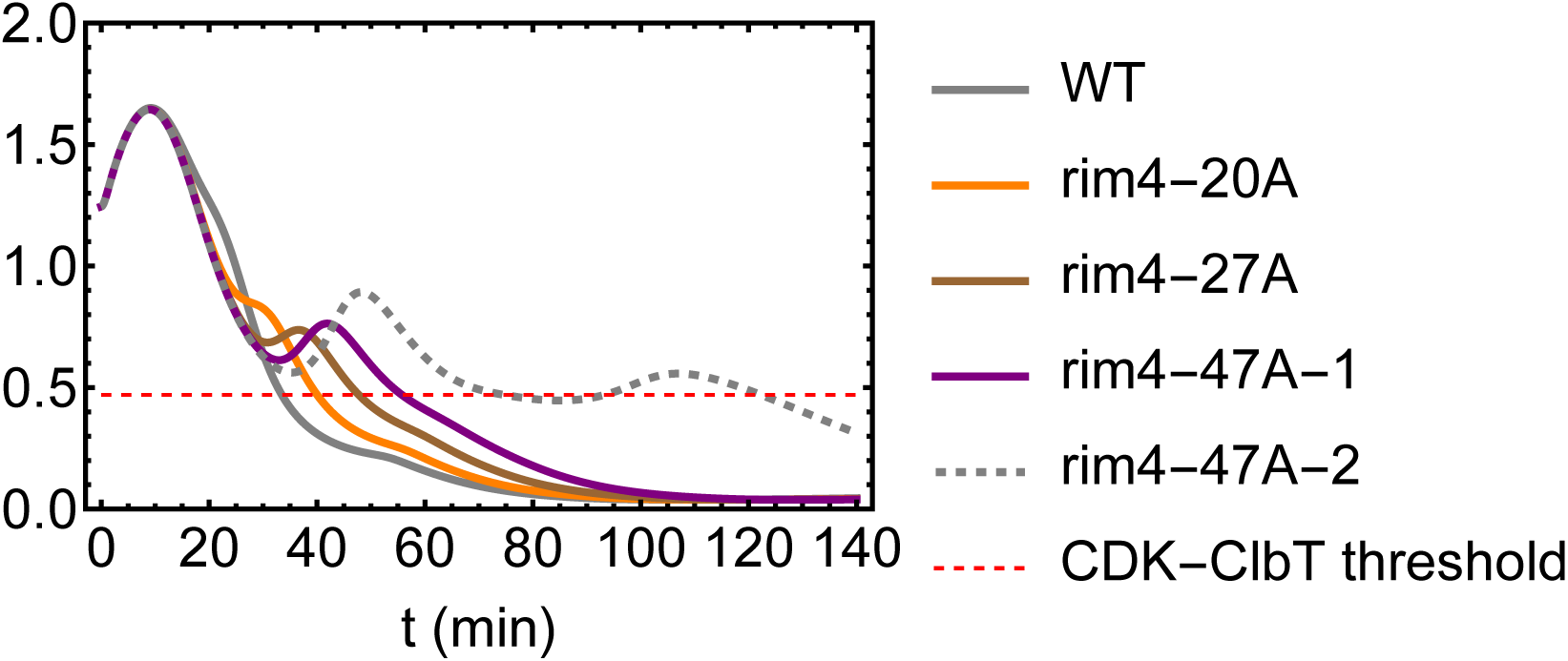
Total Cdk1-Clb concentration for WT and different Rim4 mutants. Solid gray: WT (T = 20 min, 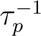 = 0.2 min^-1^). Solid orange: *rim4-20A* (T = 28 min, 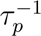 = 0.2 min^-1^). Solid brown: *rim4-27A* (T = 34 min, 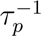 = 0.2 min^-1^). Solid purple: *rim4-47A-1* (T = 38 min, 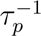 = 0.075 min^-1^). Dashed gray: *rim4-47A-2* (T = 43 min, 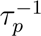 = 0.075 min^-1^). Dashed red: Cdk1-Clb concentration threshold. We note that with increasing T, and therefore delay in dissociation of mRNA from Rim4 (and Rim4 clearance), threshold crossing is delayed, until oscillatory dynamics sets in.

**Table S12:** Yeast strain list.

| **Strain number** | **Genotype** |
| --- | --- |
| LY8899 | *MATa/α, His3pr-yomRuby2-TUB1:URA3/His3pr-yomRuby2-TUB1:URA3, SPC42-GFP:HIS3/+* |
| LY9468 | *MATa/α, His3pr-yomRuby2-TUB1:URA3/His3pr-yomRuby2-TUB1:URA3, CDC14-GFP:HIS3/CDC14-GFP:HIS3* |
| LY9528 | *MATa/α, His3pr-yomRuby2-TUB1:URA3/His3pr-yomRuby2-TUB1:URA3, CDC14-GFP:HIS3/CDC14-GFP:HIS3, clb1::kanMX/clb1::kanMX* |
| LY9529 | *MATa/α, His3pr-yomRuby2-TUB1:URA3/His3pr-yomRuby2-TUB1:URA3, CDC14-GFP:HIS3/CDC14-GFP:HIS3, clb3::natMX/clb3::natMX* |
| LY9530 | *MATa/α, His3pr-yomRuby2-TUB1:URA3/His3pr-yomRuby2-TUB1:URA3, CDC14-GFP:HIS3/CDC14-GFP:HIS3, clb4::hphMX/clb4::hphMX* |
| LY9798 | *MATa/α, His3pr-yomRuby2-TUB1:URA3/His3pr-yomRuby2-TUB1:URA3, CDC14-GFP:kanMX/CDC14-GFP:kanMX, rim4-20A-3V5:HIS3/rim4-20A-3V5:HIS3* |
| LY9797 | *MATa/α, His3pr-yomRuby2-TUB1:URA3/His3pr-yomRuby2-TUB1:URA3, CDC14-GFP:kanMX/CDC14-GFP:kanMX, rim4-27A-3V5:HIS3/rim4-27A-3V5:HIS3* |
| LY9803 | *MATa/α, His3pr-yomRuby2-TUB1:URA3/His3pr-yomRuby2-TUB1:URA3, CDC14-GFP:HIS3/CDC14-GFP:HIS3, rim4-47A-3V5:kanMX/rim4-47A-3V5:kanMX* |
| LY9809 | *MATa/α, His3pr-yomRuby2-TUB1:URA3/His3pr-yomRuby2-TUB1:URA3, CDC14-GFP:HIS3/CDC14-GFP:HIS3, clb1::kanMX/clb1::kanMX, clb4::hphMX/clb4::hphMX* |
| LY9810 | *MATa/α, His3pr-yomRuby2-TUB1:URA3/His3pr-yomRuby2-TUB1:URA3, CDC14-GFP:HIS3/CDC14-GFP:HIS3, clb3::natMX/clb3::natMX, clb4::hphMX/clb4::hphMX* |
| LY9861 | *MATa/α, His3pr-yomRuby2-TUB1:URA3/His3pr-yomRuby2-TUB1:URA3, CDC14-GFP:HIS3/CDC14-GFP:HIS3, clb1::kanMX/clb1::kanMX, clb3::natMX/clb3::natMX* |
| LY9933 | *MATa/α, His3pr-yomRuby2-TUB1:URA3/His3pr-yomRuby2-TUB1:URA3, CDC14-GFP:HIS3/CDC14-GFP:HIS3, ama1::kanMX/ama1::kanMX* |
| LY10537 | *MATa/α, His3pr-yomRuby2-TUB1:URA3/His3pr-yomRuby2-TUB1:URA3, SPC42-GFP:HIS3/+, rim4-47A-3V5:kanMX/rim4-47A-3V5:kanMX* |
| LY10732 | *MATa/α, His3pr-yomRuby2-TUB1:URA3/His3pr-yomRuby2-TUB1:URA3, SPC42-GFP:HIS3/+, rim4-27A-3V5:HIS3/rim4-27A-3V5:HIS3* |
| LY10735 | *MATa/α, His3pr-yomRuby2-TUB1:URA3/His3pr-yomRuby2-TUB1:URA3, SPC42-GFP:HIS3/+, rim4-20A-3V5:HIS3/rim4-20A-3V5:HIS3* |

## Footnotes

1 Code is available publicly on github at https://github.com/abimarquez1211/Meiotic-Exit-Code.

